# Parallel clines of chromosomal inversion frequencies in seaweed flies are associated with thermal variation

**DOI:** 10.1101/2025.04.28.650981

**Authors:** Léa A. Nicolas, Emma L. Berdan, Maren Wellenreuther, Hervé Colinet, Andréa Clouard, Pierre De Wit, Sylvain Glémin, Claire Mérot

## Abstract

Chromosomal inversion supergenes, which form blocks of linked genes, are increasingly recognized for their role in maintaining intra-specific diversity. They are predicted to be relevant genetic architectures for local adaptation in the face of gene flow. However, pinpointing the underlying traits and functional mechanisms under selection remains challenging. The seaweed fly *Coelopa frigida* harbors several large polymorphic inversions, of which the *Cf-Inv(4*.*1)* inversion displays a latitudinal cline of frequencies along the North American Atlantic Coast, suggesting a putative role in adaptation along the eco-climatic gradient. To investigate this hypothesis, we designed a molecular marker for karyotyping and studied natural and experimental populations from North America and Europe. We confirmed that this inversion is also polymorphic in Europe, and displays parallel latitudinal clines across continents, providing strong indirect support that *Cf-Inv(4*.*1)* is under natural selection along similar environmental gradients. We found that *Cf-Inv(4*.*1)* had a significant impact on egg-to-adult survival and fecundity under different thermal conditions. However, no effect on cold tolerance could be determined using supercooling point and chill coma recovery time. We speculate that fitness associated with *Cf-Inv(4*.*1)* is shaped by subtle life-history differences whose relative advantage depends on climate. While our experimental approaches provided insights into genotype-phenotype associations, it is worth noting that selection acts on the overall fitness, involving complex sets of traits. This is especially relevant for inversions linking hundreds of genes. This multi-gene property also explains why inversions are frequently involved in repeated parallel adaptation to environmental gradients, as demonstrated here in the seaweed fly.

## Introduction

From the early discovery of chromosomal rearrangements by cytogenetics to the recent advances in whole genome sequencing, an increasing body of evidence has demonstrated the importance of structural genomic variation in adaptation and diversification. A chromosomal inversion is a structural variant defined as a segment of a chromosome in which two sequences in reversed orientations segregate in the population (Allendorf et al., 2012). This different orientation translates into reduced gene flow between the two arrangements (the standard and the inverted sequence), as recombination is limited in heterokaryotypes (Tigano & Friesen, 2016). Therefore, the two arrangements tend to diverge through time and, due to high linkage disequilibrium, can behave as two alleles of one single gene. As inversions can link multiple genes that jointly contribute to complex phenotypes, these inversions can be referred to as “supergenes”. This type of polymorphism is often associated with speciation, the specialization of ecotypes, and local adaptation (Wellenreuther & Bernatchez, 2018), as chromosomal inversions can preserve combinations of locally-adapted alleles, thereby limiting the effect of outbreeding depression in contexts of high gene flow (Kirkpatrick & Barton, 2006; Schaal et al., 2022, Yeaman, 2013). Owing to this property, and because inversion polymorphisms can be retained over large spatial and temporal scales, chromosomal inversions are also predicted to be repeatedly involved in parallel adaptation to similar ecological gradients (Westram et al., 2022).

The importance of chromosomal inversions in local adaptation and intra-specific diversification is supported by empirical cases in which adaptive ecotypes and morphotypes have been associated to inversion polymorphisms in plants, insects, birds, and mammals (Hager et al., 2022; Jay et al., 2022; Lindtke et al., 2017; Purcell & Brelsford, 2025; Wellenreuther & Bernatchez, 2018). Besides the charismatic cases in which the inversion-phenotype associations have been well-described, more and more studies are also detecting large polymorphic inversions in genome scans (Fang & Edwards, 2024; Li et al., 2023; Liang et al., 2024). However, understanding the functional and putatively adaptive effects of those inversions remains a challenge. Clues about the selective forces acting on inversions can be inferred from the distribution of the polymorphism in natural populations, particularly when parallel patterns are observed across similar environmental gradients. For example, sharp frequency changes in multiple inversions in *Littorina saxatilis*, repeatedly observed across a habitat transition, support their role in local adaptation (Faria, Chaube, et al., 2019). Similarly, chromosomal inversions in *Drosophila melanogaster* form parallel frequency clines across continents and seasons, suggesting that the inversion polymorphisms are under selection by an environment that varies in space and time (Kapun & Flatt, 2019). In *Calosoma* beetles, several chromosomal inversions overlap with regions of high divergence between highland and lowland ecotypes in a context of repeated highland adaptation in the Galápagos (Vangestel et al., 2024). Such observational data benefit from further experiments. For instance, in *Drosophila subobscura*, inversions were associated with different responses to thermal stress in the laboratory, partially explaining the observed latitudinal clines in frequencies (Rego et al., 2010). The clinal inversion polymorphism In(3R)P in *Drosophila melanogaster* contributes to latitudinal variation in multiple fitness-related traits, acting as a ‘supergene’ likely maintained by fitness trade-offs and balancing selection across geographic regions (Durmaz et al., 2018). Overall, field-based and lab-based studies of inversion polymorphisms thus remain critical to understand the functional impact of the multiple inversions currently being detected by genomic tools. Only by elucidating the relationship between inversion, fitness, and environment can we understand the spatial and temporal dynamics of those large variants and the evolution of their genomic content.

In the seaweed fly *Coelopa frigida*, four large polymorphic inversions have been detected in North American populations (Mérot et al., 2021). One of them, located on chromosome 1, *Cf-Inv(1)*, which is known to impact adult size, development time, and a set of life-history traits (Butlin et al., 1982; Butlin & Day, 1984; Mérot et al., 2020) exhibits patterns of local adaptation (Wellenreuther et al., 2017). However, the phenotypes associated with the three inversions located on chromosome 4 (*Cf-Inv(4*.*1), Cf-Inv(4*.*2), Cf-Inv(4*.*3)*) remain unknown. Interestingly, their frequencies co-vary with latitude along the North American East Coast (Mérot et al., 2021), particularly strongly for *Cf-Inv(4*.*1)*, which ranges from 25% to 75% over a 1000km North-South gradient. Such a latitudinal cline suggests that *Cf-Inv(4*.*1)* may play a role in climate adaptation, possibly in thermal adaptation. Indeed, temperature is one of the main environmental factors varying with latitude and is expected to be a strong selective force for ectotherm insects (Denlinger & Lee Jr, 2010), especially for *C. frigida* whose distribution extends from temperate to polar regions (Dobson, 1974). Cytogenetic studies (Aziz, 1975) have suggested that the inversion *Cf-Inv(4*.*1)* may also be carried by European populations, providing an ideal system to assess the contribution of chromosomal inversions to parallel adaptation along ecogeographic gradients, as well as providing natural replicates to investigate the putative functional impact of the inversion. However, differences in the karyotype frequency along the latitudinal gradient in Europe have yet to be investigated.

Here, we tested whether the genomically identified North American inversion *Cf-Inv(4*.*1)* could also be detected in European populations of *C. frigida*. After validating a genetic marker to genotype the inversion on both continents, we investigated the geographic distribution of frequencies along a North-South cline in Europe. The demonstration of inter-continental parallelism would provide indirect evidence that the latitudinal clines of inversion frequencies are the result of divergent selection based on latitude, and not the simple effect of neutral processes such as genetic drift and isolation by distance (Westram et al., 2022). Through an experimental approach, we then tested the hypothesis that the cline of inversion frequencies reflects the role of *Cf-Inv(4*.*1)* in adaptation to the latitudinal thermal gradient. We tested the effect of *Cf-Inv(4*.*1)* on cold tolerance (supercooling point, chill coma recovery time) and measured fitness components (fecundity, longevity, development time, egg-to-adult survival) under different thermal conditions for both inversion homokaryotypes.

## Methods

### Study organism and field sampling

*Coelopa frigida* is an acalyprate fly (Diptera: Coelopidae) that lives and feeds in the banks of decomposing seaweeds on the shores of the Atlantic Ocean. It is distributed from Massachusetts (USA) to the Canadian Arctic in North America and from Brittany (France) to Svalbard (Norway) in Europe.

Adult flies were collected along the coasts of the Atlantic Ocean in six North American locations (Supplementary Table 1) during summer 2022 and ten European locations during spring 2023 (Fig. 1A, Tab. 1). Samples were either stored in ethanol 96% or stored at -80°, while 100-300 individuals per population were kept alive to start 6 laboratory populations per continent. Those were subsequently maintained for several generations in controlled laboratory conditions (see details in Supplementary Methods 1).

**Table 1.**
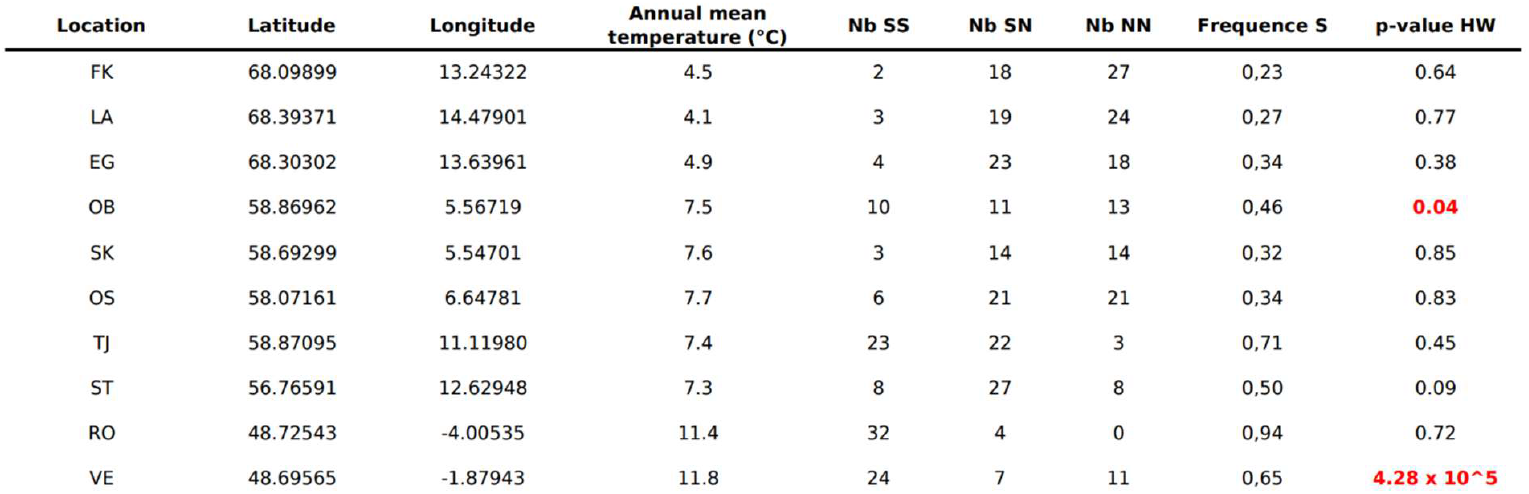
Table of European locations, coordinates, mean annual air temperature, karyotypes counts, frequency of arrangement S, and p-values of the Chi-squared test of Hardy-Weinberg equilibrium.

| Location | Latitude | Longitude | Annual mean temperature (°C) | Nb SS | Nb SN | Nb NN | Frequency S | p-value HW |
| --- | --- | --- | --- | --- | --- | --- | --- | --- |
| FK | 68.09899 | 13.24322 | 4.5 | 2 | 18 | 27 | 0,23 | 0.64 |
| LA | 68.39371 | 14.47901 | 4.1 | 3 | 19 | 24 | 0,27 | 0.77 |
| EG | 68.30302 | 13.63961 | 4.9 | 4 | 23 | 18 | 0,34 | 0.38 |
| OB | 58.86962 | 5.56719 | 7.5 | 10 | 11 | 13 | 0,46 | **0.04** |
| SK | 58.69299 | 5.54701 | 7.6 | 3 | 14 | 14 | 0,32 | 0.85 |
| OS | 58.07161 | 6.64781 | 7.7 | 6 | 21 | 21 | 0,34 | 0.83 |
| TJ | 58.87095 | 11.11980 | 7.4 | 23 | 22 | 3 | 0,71 | 0.45 |
| ST | 56.76591 | 12.62948 | 7.3 | 8 | 27 | 8 | 0,50 | 0.09 |
| RO | 48.72543 | -4.00535 | 11.4 | 32 | 4 | 0 | 0,94 | 0.72 |
| VE | 48.69565 | -1.87943 | 11.8 | 24 | 7 | 11 | 0,65 | **4.28 x 10<sup>-5</sup>** |

**Fig. 1.**
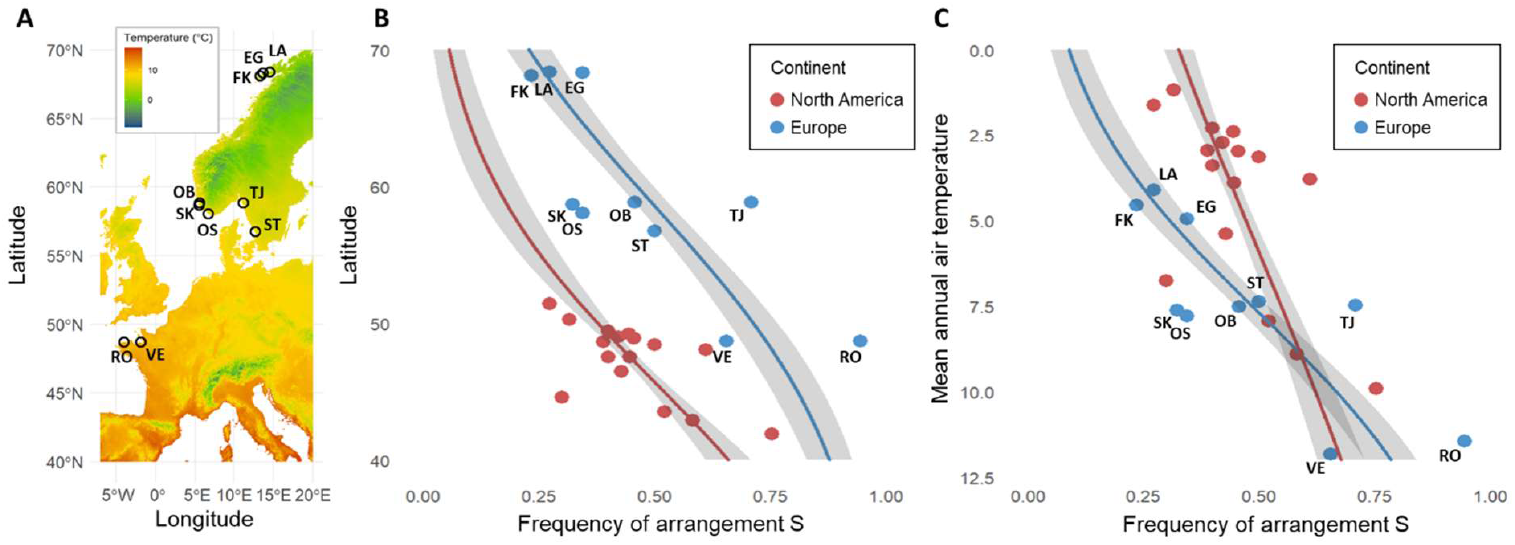
*Coelopa frigida* sampling and inversion frequency clines along the latitudinal gradients. (A) Map of sampling locations in Europe. The color gradient represents mean annual air temperatures. (B) Frequency of *Cf-Inv(4*.*1)* arrangement S as a function of latitude (C). Frequency of *Cf-Inv(4*.*1)* arrangement S as a function of mean annual air temperature. Red dots represent the North American data provided by Mérot *et al*. (2021) and blue dots represent the European data, from 10 locations sampled in this study. Lines are the binomial regressions and the grey ribbons are the confidence interval 95% of the average predictions of the binomial model, which accounts for sample sizes.

### Detecting inversion *Cf-Inv(4*.*1)* in Europe

The inversion *Cf-Inv(4*.*1)* was inferred by principal component analysis (PCA) on low-coverage whole genome sequences from 1,446 North American *C. frigida* (Mérot et al., 2021). Its position ranges approximately from position 1,088,816 to position 7,995,568 on chromosome 4 based on recombination reduction, and has two main arrangements, hereafter named S (South) and N (North). To test whether the inversion was also present and polymorphic in Europe, we used a dataset of 56 South Scandinavian *C. frigida* collected and sequenced in 2015 with paired-end short-reads 150 bp in an Illumina Hi-Seq machine at a coverage of ∼30X at SciLifeLab (Sweden). We completed this dataset with a subset of 104 individuals extracted from the North American low-coverage (1X) whole genome sequences used in Mérot et al. (2021). Reads were trimmed and aligned with SnpArcher (Mirchandani et al. (2024), accessed on 3 Apr 2025) to the reference genome of *Coelopa frigida* (NCBI accession: PRJNA688905, Mérot et al, 2021). To account for depth heterogeneity, genotype likelihoods and *F*_ST_ statistics were calculated with ANGSD v0.940 (Korneliussen et al., (2014), accessed on 23 Apr 2025). We used Winpca v 1.2 (Blumer et al., 2025) and pcangsd (Meisner & Albrechtsen, 2018) to perform a sliding-window PCA along chromosome 4. Based on this analysis, the putative inversion position was restricted to the interval 1,136,457-5,099,894. In this region, we performed a PCA and visually assigned putative karyotypes for the inversion *Cf-Inv(4*.*1)* based on the three observed clusters.

To investigate the differentiation within and outside the inversion region across continents and arrangement, we then calculated *F*_ST_ as pairwise comparisons across the four groups: SS-America, SS-Europe, NN-America, NN-Europe. For each pair, we calculated *F*_ST_ inside the inversion, in chromosome 4, and in sliding-windows of 25 kb.

### Developing a diagnostic marker to genotype the inversion

We aimed to develop a SNP marker which would allow us to genotype with a simple PCR and digestion assay. The marker was first developed based on the North American WGS dataset. Several SNPs were selected according to the following conditions: (1) located in coding regions, (2) fixed between the two arrangements (at least a difference of allele frequency of 99% between the two homokaryotypic groups), (3) alternatively cut or not cut by a common restriction enzyme. After several assays, we selected two linked SNPs (located on chromosome 4 at position 1,637,841 and 1,637,843) in a 667 bp-long coding region, corresponding to the gene DN9360 (chr4:1,636,902-1,638,332), which amplified consistently well (see primers in Supplementary Table 2) and where the restriction enzyme EcoRI cut the arrangement S but not the arrangement N. The whole protocol of the diagnostic assay is available in supplementary materials (Supplementary Methods 3 and 4; Supplementary Figure 1). We performed the diagnostic assay using the DNA of the same subset of 104 North American individuals used for the PCA (see DNA extraction method in Mérot et al., 2021) and calculated the concordance between karyotypes determined with the marker and those given by the whole-genome PCA. We used the same approach to validate the marker in Europe, using the DNA extracts provided by Emma Berdan (see Supplementary Methods 2 for extraction protocol) for a subset of 38 samples out of the 56 used for the PCA (no DNA could be recovered from the remaining samples). The amplified fragment of 667 bp, was also sequenced at the Plate-forme d’Analyses Génomiques (Université Laval, Québec, Canada) for 8 North American samples and at Macrogen Europe for 13 Scandinavian samples. The forward and reverse sequences were processed using Geneious 2023.0.4, aligned by pairs, trimmed at 488 bp, corrected for sequencing ambiguities, and realigned altogether. To assess sequence variation within and across continents, a haplotype network was built using the Popart software (http://popart.otago.ac.nz) through the Minimum spanning network method (Bandelt et al., 1999) with ɛ = 0.

### Investigating the distribution of inversion frequencies in Europe

In the 10 European locations spanning a latitude from 48.69565° to 69.39371° (Fig. 1A), we used the diagnostic marker to genotype 31-48 flies per location.

All statistical analyses, described in this section and further, were performed on R version 4.4.3. The frequencies of the two arrangements were calculated for each location, as well as the proportion of the three karyotypes (SS, SN, NN). Hardy-Weinberg equilibrium in the karyotypes proportion was tested for using the function HWAlltests from the package *Hardy-Weinberg*. To assess the correlation between inversion frequencies and a putative climatic cline, annual mean temperature data were extracted from the database ‘Worldclim’ (Fick & Hijmans, 2017) and the associations between frequencies and latitude/temperature were tested with a binomial regression using the glm function with the link “logit”.

### Experimentally testing the role of inversion *Cf-Inv(4*.*1)* in response to cold stress

To test the predictions that cold tolerance varies across locations along the latitudinal cline and is associated with inversion karyotypes, we experimentally measured two traits reflecting cold stress tolerance: the supercooling point (SCP) and the chill coma recovery time (CCRT). Experiments were performed on 288 and 320 individuals respectively, from five North American locations (see Supplementary Table 1) raised in lab for 3-8 generations. Detailed methods are available in supplementary materials (Supplementary Methods 5 and 6). Briefly, the supercooling point was defined as the temperature at the onset of the freezing exotherm peak for each fly exposed to a gradually decreasing temperature at a rate of -0.5°C/min (from 5 to -25°C). Chill coma recovery time was defined as the time before the insect spontaneously recovers coordinated neuromuscular function and stand on legs at 15°C, following a prolonged chill coma induced by an exposure at -5°C for 16h (equivalent to a cold winter night). Experimental individuals were genotyped for inversion *Cf-Inv(4*.*1)* using the protocol described above (Supplementary Methods 3 and 4). We also controlled for the inversion *Cf-Inv(1)* by genotyping following a protocol described in Mérot et al (2018). Supercooling point was normally distributed and analyzed using a Linear Model (function lm). Chill coma recovery time was analyzed with a Generalized Linear Model (GLM) with Gamma family, which is appropriate for a time-to-event variable. Both models included mass, sex, location, and karyotypes for both inversions as explanatory variables, and were tested with the function Anova (type II) from the R package *car* (Fox & Weisberg, 2018), which performs an F test on LMs and a Likelihood Ratio Test on GLMs. When necessary, pairwise permutation t-tests or log-rank tests were performed to identify the modalities that differed from others.

### Experimentally testing the combined effects of inversion *Cf-Inv(4.1)* and temperature on fitness

#### Creating homokaryotypic lineages for *Cf-Inv(4.1)*

To perform fitness experiments, we made four experimental lines (North American SS and NN, European SS and NN) which were homokaryotypic for the marker of the inversion *Cf-Inv(4*.*1)* but highly variable in the rest of the genome. To start the 1^st^ generation, 700 virgin individuals from the laboratory populations were genotyped with the diagnostic marker (see Supplementary Methods 3 and 4), using a piece of leg cut under CO_2_ anesthesia. A total of 314 homokaryote individuals were selected for egg laying (N=43 for SS-America, N=92 for NN-America, N=24 for SS-Europe, N=155 for NN-Europe). Those lines were maintained at 20°C (reference temperature) using the raising method described in supplementary materials for six generations before starting the experiments. The karyotype of 12 randomly chosen individuals per lineage was checked every two generations, by genotyping them for *Cf-Inv(4*.*1)* with the diagnostic marker.

#### Measuring fitness during larval development

Larval fitness was assessed by two traits: development time and viability (i.e., survival from egg to adult). For each of the four homokaryote lines, we collected a large number of eggs which were randomly divided, using a paint brush soaked in sea water, into 30 batches of approximately 100 eggs. Each batch was spread on a piece of *Saccharina latissima* and photographed to be counted *a posteriori* using ImageJ 1.54b (Supplementary Figure 9) before being placed in a plastic box of dimension 10.5 × 7.5 × 11 cm filled with 90 g of seaweeds (50% *Fucus sp*., 50% *Saccharina sp*.). The 30 boxes were distributed randomly across five thermal conditions: 10, 15, 20, 25 and 30°C (six batches of ∼100 eggs per temperature per lineage). A total of 45 g of algae was added at half of the development time, i.e., on day 3, 5, 7, 14 and 31 for temperatures 30, 25, 20, 15 and 10°C respectively. During the emergence phase, eclosed adults were removed from the boxes daily, counted and sexed. Data at 10°C were not statistically analyzed because of a very low viability (< 5%), which resulted in a very low sample size (n = 4).

#### Measuring fitness during the adult stage

Adult fitness was assessed by measuring two life-history traits: adult longevity (*i*.*e*., time between adult emergence and death) and female fecundity. For each of the four homokaryote lines, we used 36 virgin individuals to create 18 couples, that were split across three temperatures: 10, 20 and 30°C (six couples per temperature per line). To ensure virginity, individuals were collected at the pupal stage, kept individually at 20°C, and sexed upon emergence. Because females generally emerge a few days before males, they were first placed alone in plastic boxes of dimensions 8 × 8 × 5 cm with a piece of *Saccharina latissima* and a piece of *Fucus vesiculosus* of approximately 3 × 3 cm, and split across the three experimental conditions. A male from the same line was added one to two days later. From the following day, the boxes were checked on a regular basis (every day at 20°C and 30°C, every other day at 10°C) to count clutches of eggs, replace the pieces of seaweed, and check for dead individuals. The death of a female meant the end of the fecundity measurement but the box was kept at the experimental temperature to continue measuring the remaining male’s longevity. Dead males were replaced by new males of the same lineage (whose longevity was not measured). To estimate female fecundity, we measured: (i) the size of the first clutch, (ii) the mean clutch size, (iii) the total number of eggs laid, and (iv) the number of clutches laid during the entire life.

#### Disentangling possible confounding factor effects from *Cf-Inv(1)*

Besides *Cf-Inv(4*.*1), C. frigida* carries another polymorphic inversion on chromosome 1, *Cf-Inv(1)*, which affect life-history traits such as development time, survival, and fecundity. We controlled for the frequency of *Cf-Inv(1)* in the experimental lines by genotyping a subsample of 35 to 48 individuals per line (see methods in Mérot et al., 2018). The frequency of *Cf-Inv(1)* was analyzed with a Chi-square test of homogeneity.

#### Statistical analyses

Variation in each response variable (longevity, female fecundity, development time, viability) was analyzed with GLMs including the following explanatory variables: inversion karyotype, temperature and continent. For longevity and development time, we also tested the effect of sex. The linear models were implemented in R using the package *stats* (R Core Team, 2025). Development time and longevity were analyzed using a GLM based on a Gamma distribution, being time-to-event variables. Viability data were analyzed using binomial GLM with a logit link function for binary outcome (dead vs. alive). Number of clutches and total fecundity, expected to follow a Poisson distribution, were analyzed with a GLM based on a Poisson, or a Quasi-Poisson law to address the overdispersion of residuals. First clutch size and mean clutch size, expected to follow a normal distribution, were analyzed with a GLM with quasi family (link = identity) because of overdispersion of the residuals with a simple linear regression. Accounting for the relatively small sample size, especially for viability and female fecundity, we decided to remove interactions of more than two factors.

After validating independence of the residuals, we investigated the explanatory contribution of each variable with deviance analyses (function Anova from the package *car* (Fox & Weisberg, 2018)). When necessary, multiple comparisons were performed using the method of Estimated Marginal Means (function emmeans from the package *emmeans* in version 1.8.8 (Lenth *et al*., 2023)), to test the difference between modalities of significant factors and between combinations of modalities in the case of interactions. R^2^ was estimated based on model deviance, as 1 – (model deviance / null model deviance).

We modelled the thermal performance curves for viability with a simple quadratic model (*viability* = *a* × *temperature*^2^ + *b* × *temperature* + *c*), using R packages *rTPC* and *nls_multstart* (Padfield et al., 2021). We extracted the estimated values of maximal performance (rmax), optimal temperature (Topt), critical thermal limits (CTmin and CTmax), thermal tolerance (amplitude of temperature for which viability is above zero), and breadth (amplitude of temperature for which viability is above 80% of rmax).

## Results

### The inversion *Cf-Inv(4*.*1)* is shared across continents

Whole genome sequencing of European *C. frigida* supported the presence of a polymorphic inversion in the same genomic region as *Cf-Inv(4*.*1)*. On PCAs performed on small windows of 50 kb, the first principal component decomposing genetic variation separated individuals by continents along most of chromosome 4, except within the inversion putative boundaries, where individuals formed three distinct clusters including samples from both continents, corresponding to the three karyotypes of *Cf-Inv(4*.*1)* (Fig. 2A). The sliding-window PCA also revealed that this low-recombing region associated with the inversion is actually shorter than estimated in Mérot et al. (2021), particularly in Europe. Genetic variance within the newly estimated inversion boundaries (chr4:1,136,457-5,099,894) is structured in three clusters along the first PC (explaining 32.02% of variance), corresponding to the three karyotypes of *Cf-Inv(4*.*1)* in North America, and in two groups along the second PC (explaining 5.86% of variance) corresponding to continents (Fig. 2B).

**Fig. 2.**
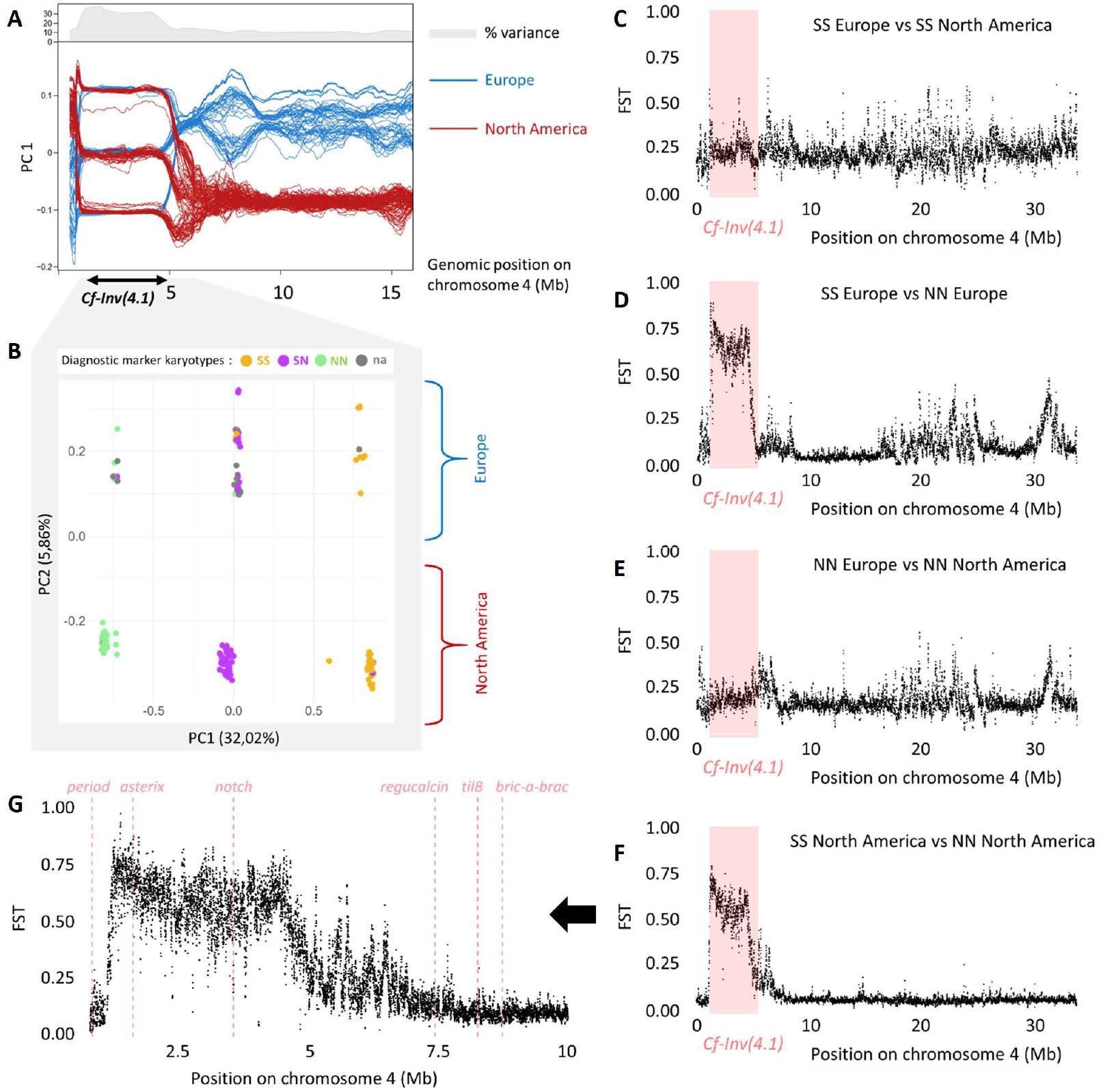
Inversion *Cf-inv(4*.*1)* is shared across continents. **(A)** Sliding-window PCA decomposing genetic variation captured by SNPs along chromosome 4. Red lines represent the PC1 score of North American individuals. Blue lines represent the PC1 score of European individuals. Grey zone on top corresponds to the percentage of variance explained by PC1. **(B)** PCA decomposing genetic variation captured by SNPs between positions 1,136,457 and 5,099,894, corresponding to the low-recombining region associated with inversion *Cf-inv(4*.*1)* as determined graphically on the sliding-window PCA. PC1 explains 32,02% of the variance and displays 3 clusters, characteristic of a non-recombining region. PC2 explains 5,86% of variance. Points are colored according to the karyotypes given by the designed diagnostic marker. Grey points represent the samples for which no DNA was available for amplification. **(C, D, E, F)** FST along chromosome 4, comparing **(C)** European SS versus North America SS individuals, **(D)** European SS versus European NN individuals, **(E)** European NN versus North American NN individuals, **(F)** North American SS versus North American NN individuals. Pink zone correspond to the location of *Cf-Inv(4*.*1)* detected in North America. **(G)** Zoom on FST variation from positions 0.8 to 10 Mb comparing SS versus NN in North America. Purple lines represent the positions of the candidate genes.

Although genetic differentiation between continents was non-negligeable, with *F*_ST_ around 0.13 across the genome, this continental differentiation was higher within each arrangement of *Cf-Inv(4*.*1)* (0.20 among NN individuals, 0.22 among SS individuals, Fig. 2C-E). In sharp contrast, *F*_ST_ between the arrangements (NN vs SS), and within the inversion boundaries, was very high on both continents (0.62 in North America, 0.66 in Europe, Fig. 2D-F). Such a pattern of high differentiation between arrangements along *Cf-Inv(4*.*1)* and low differentiation between continents is also observed in the coding fragment including the proposed diagnostic marker (Fig. 3). The sequences of arrangements N and S differed by a minimum of 16 nucleotides (3.3% divergence), while the difference between North American and European sequences of the same arrangement was either one or two nucleotides (0.4% divergence).

**Fig. 3.**
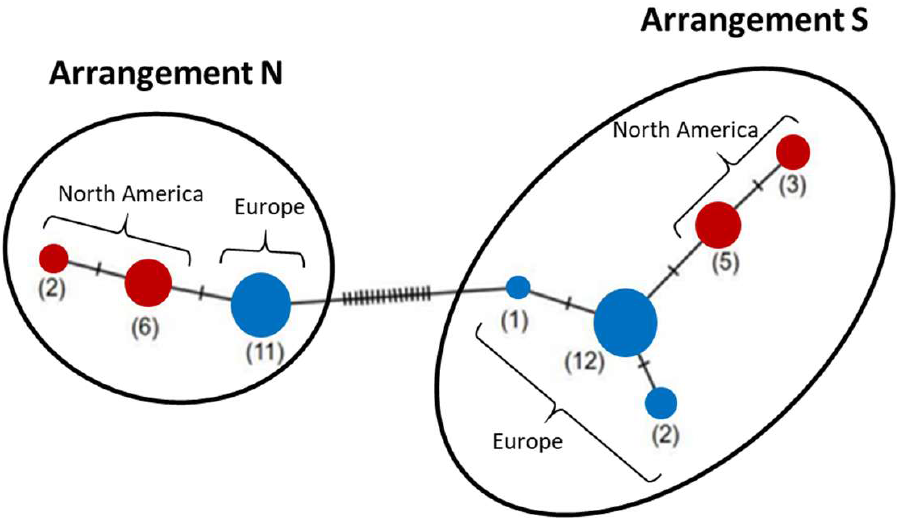
Haplotype network representing similarities and differences between haplotypes from 8 North American and 13 Scandinavian individuals at a region of 488 pb around the diagnostic marker. Circle areas are proportional to the number of haplotypes with identical sequences. Sticks represent the number of substitutions between two circles. Two main haplotype groups were found corresponding to the S and the N arrangements, as labelled. The network was built with the Minimum spanning network method.

This high sequence differentiation across arrangement but similarity across continents provided the opportunity to develop a simple diagnostic marker. In North America, the diagnostic marker showed a very good accuracy with 99% concordance with inversion karyotypes determined by PCA (Mérot et al., 2021). Out of 103 individuals genotyped with the marker, 102 were consistent with the karyotype obtained from the PCA on whole genome sequences. In Europe, 32 out of 38 individuals were consistent, representing an operational concordance rate of 84% (sees details in Supplementary Table 3). To examine the cause of these inconsistencies, we called the genotypes of all SNPs within a region of 6 kb around the two diagnostic SNPs (see Supplementary Figure 3 and Supplementary Results 4). The SNPs were consistent with the PCA in 1 out of 1 in North America and 2 out of 6 in Europe, suggesting that 1 to 5% of the discordance may be due to technical issues such as gel reading/enzymatic digestion. In Europe, the remaining 4 individuals presented a short track of SNP genotypes incoherent with the karyotypes inferred from PCA, possibly revealing a higher shared polymorphism across the S and N arrangements in Europe and/or the result of gene conversion.

### Inversion *Cf-Inv(4*.*1)* forms two parallel latitudinal clines

The inversion *Cf-Inv(4*.*1)* showed a cline of frequencies co-varying with latitude and annual mean air temperature in Europe (Fig. 1), as in North America (Mérot et al., 2021). The relationship between latitude and the frequency of the arrangement S is significant on both continents with a slope of -0.12 (binomial regression, χ^2^ = 67.66, df = 1, p-value < 0.001, R^2^ = 0.82) for North America and a slope of -0.11 (binomial regression, χ^2^ = 100.44, df = 1, p-value < 0.001, R^2^ = 0.93) for Europe. It is worth noting that the association between latitude and inversion frequency in Europe also holds true when using a higher-confidence marker, *i*.*e*. only individuals genotyped by PCA (binomial regression, χ^2^ = 8.18, df = 1, p-value = 0.004, Supplementary Figure 4A). We also found a significant relationship between the inversion frequency and the annual mean temperature in both North America, with a slope of 0.12 (Fig. 1C, binomial regression, χ^2^ = 69.93, df = 1, p-value < 0.001, R^2^ = 0.83) and in Europe, with a slope of 0.30 (Fig. 1C, binomial regression, χ^2^ = 101.51, df = 1, p-value < 0.001, R^2^ = 0.93). The association between inversion frequency and with minimal winter temperature gave consistent results (see Supplementary Figure 5). All European locations were at the Hardy-Weinberg equilibrium (Table 1, Chi-squared test, p-value > 0.05), except OB (58.9°N, 5.6°E), in which heterokaryotypes were slightly under-represented (χ^2^ = 4.11, p-value = 0.043); and VE (48.7°N, 1.9°W), in which heterokaryotypes were strongly under-represented (χ^2^ = 16.74, p-value < 0.001). In North America, only one location (KA, *cf*. Mérot et al., 2021) was not at the Hardy-Weinberg equilibrium (χ^2^ = 4.55, p-value = 0.033), with heterokaryotypes being slightly over-represented.

### Coelopa frigida exhibits a high cold tolerance, not associated with Cf-Inv(4.1)

Cold tolerance experiments demonstrated that *C. frigida* has a low freezing temperature, with a mean supercooling point (SCP) of -16°C (CI95% = [-16.4; - 15.6]), typical of freeze-avoidant species. The CCRT experiment also suggested a high chilling tolerance as after 16h at -5°C, only 2.8% of *C. frigida* were dead while 97.2% recovered. Recovery was also relatively fast with a recovery time ranging from 4.5 minutes to 26.5 minutes, and a mean value of 12.2 minutes (CI95% = [-11.8; 12.6]). Neither the SCP nor the CCRT were significantly affected by the karyotype at inversion *Cf-Inv(4*.*1)* despite testing a total of 65 SS, 104 SN and 97 NN for SCP (F = 0.98, df = 2, p-value = 0.38) and 41 SS, 113 SN and 89 NN for CCRT (χ^2^ = 4.67, df = 2, p = 0.097). Small effects associated with mass and sex, as well as location and karyotype at inversion *Cf-Inv(1)* were detected but remained marginal in explaining the total variance (see Supplementary Results 1).

### Egg-to-adult viability is impacted by temperature and *Cf-Inv(4*.*1)*

Temperature had a significant impact on the viability of the developing individuals (χ^2^ = 68.48, df = 3, p-value < 0.001, Fig. 4). Maximum viability was at 20°C (0.26 ± 0.14), and decreased as temperature deviated from this optimum both towards lower or higher temperatures. At 10°C, viability was very low and adult flies were recovered in only two replicates out of 24 replicates but always in NN (one European NN with 2.5% of viability and one North American NN with 4% of viability). Karyotype (χ^2^ = 12.60, df = 1, p-value < 0.001), continent (χ^2^ = 14.07, df = 1, p-value < 0.001), and their interaction (χ^2^ = 174.05, df = 1, p-value < 0.001) significantly impacted viability (Fig. 4). When considering all thermal conditions together SS karyotypes had an overall higher viability than NN karyotypes (χ^2^ = 12.6, df = 1, p-value = < 0.001) although this result hides differences across continents. Viability was significantly higher for SS than NN in North America (z = 10.80, p-value < 0.001), whereas the opposite effect was observed in Europe (z = 6.97, p-value < 0.001). We also detected a significant effect of the interaction between inversion karyotype and temperature (χ^2^ = 35.27, df = 3, p-value < 0.001), reflected in the slightly different shape and width of thermal performance curves (Supplementary Figure 6). Thermal curves revealed higher maximal viability and a higher amplitude of tolerated temperatures for the SS karyotype over NN in North America, but the opposite effect in Europe (Supplementary Table 4).

**Fig. 4.**
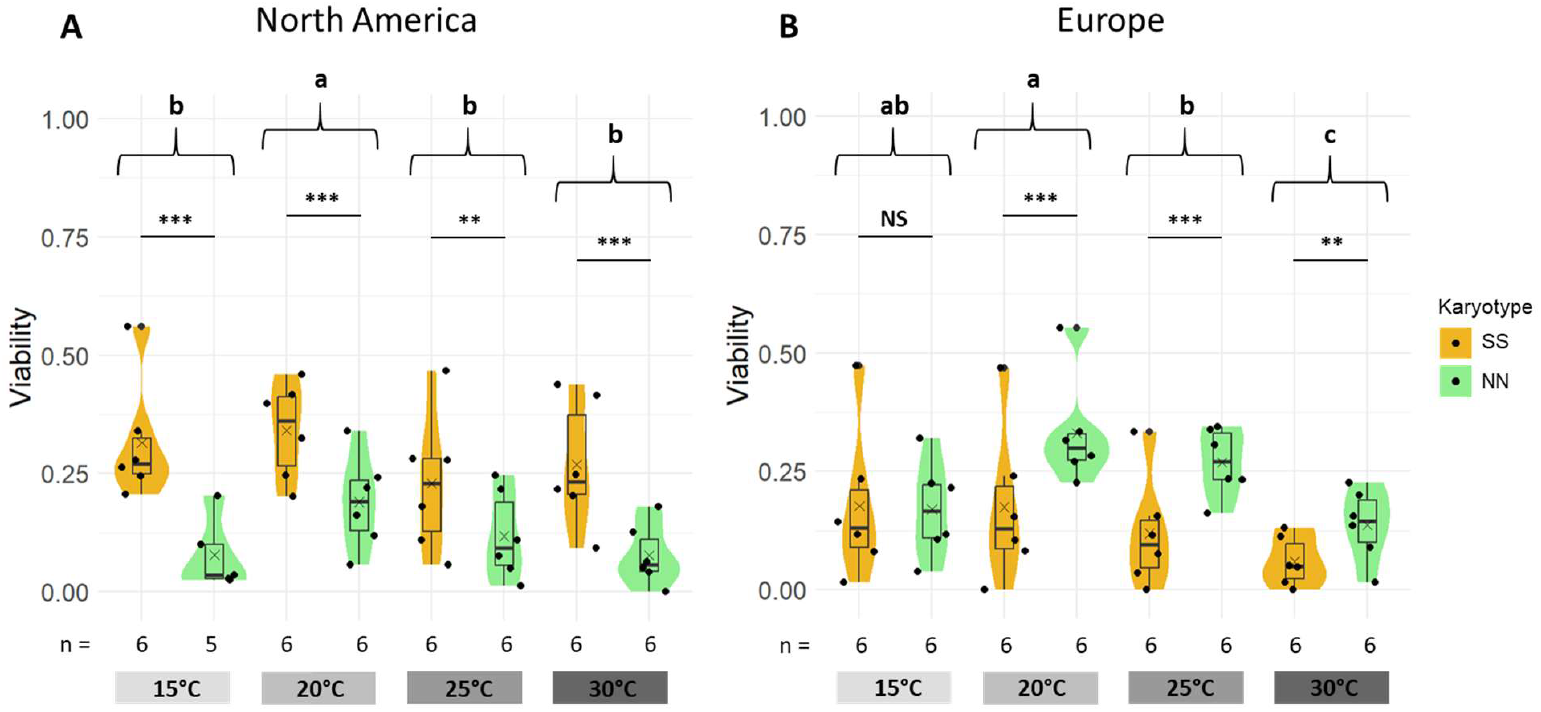
Egg-to adult viability. Viability for each combination of temperature and inversion karyotype in North America (A) and in Europe (B). Viability was quantified as the percentage of eggs developing until emerging adults under controlled temperature (10, 15, 20, 25 and 30°C). Data are not shown for 10°C because only two replicates produced adults with a very low survival rate (<5%).

### Development time is mostly determined by sex and temperature

Development time was mostly influenced by temperature (χ^2^ = 49 566, df = 3, p-value < 0.001), as the model including temperature alone explained 96.6% of the variance. Development time decreased significantly when the temperature increased (8.1 ± 0.9 days at 30°C; 9.9 ± 1.1 days at 25°C; 15.2 ± 1.2 days at 20°C; 28.1 ± 2.1 days at 15°C; 87 ± 10.4 days at 10°C; Supplementary Figure 7). Across all thermal conditions, females developed faster on average than males (χ^2^ between 97 and 250, df = 1, p < 0.001 at all temperatures), with males emerging on average 1, 1.1, 1.5 and 2.2 days after the females at 30, 25, 20 and 15°C respectively. Generally, the inversion *Cf-Inv(4*.*1)* did not have any significant effect on development time (χ^2^ = 0, df = 1, p-value = 0.55) although we detected a very small effect size at 15°C interacting with sex (χ^2^ = 11.01, df = 1, p < 0.001) and continent (χ^2^ = 10.65, df = 1, p < 0.01).

### Adult longevity is mostly determined by temperature

Analysis of adult longevity revealed no effect of *Cf-Inv(4*.*1)* but a strong effect of temperature, (χ ^2^ = 799, df = 2, p-value < 0.001, Fig 5C). The longevity at 10°C (34 ± 14 days) was higher than the longevity at 20°C ((11 ± 4 days, t = 14.02, df = 108, p-value < 0.001), which was itself higher than at 30°C (5 ± 1 days, t = 9.51, df = 108, p-value < 0.001). Continent and sex also significantly affected longevity (continent: χ ^2^ = 16.46, df = 1, p-value < 0.001; sex: χ ^2^ = 8.06, df = 1, p-value < 0.01), but post-hoc pairwise comparisons suggest a very marginal effect (with females living slightly longer than males, and North American flies living slightly longer than European flies). Adult longevity did not differ significantly between inversion karyotypes (χ ^2^ = 0.7, df = 1, p-value = 0.4, Fig 5C), even when comparing karyotypes within temperature modalities. The global model explained 87% of variance in longevity.

**Fig. 5.**
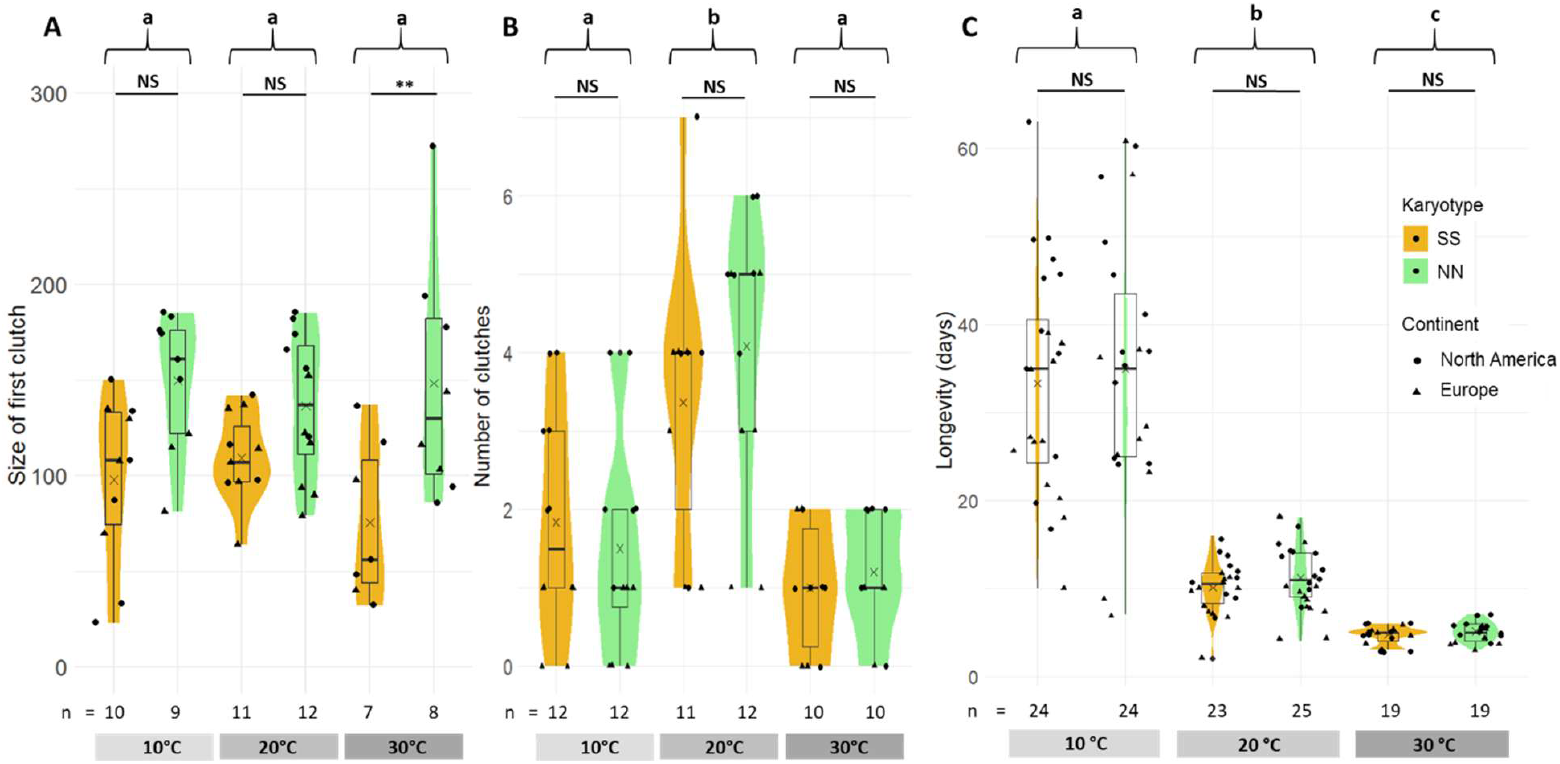
Fitness at adult stage. (A) Number of eggs laid in the first clutch of females for each combination of continent and karyotype. (B) Number of clutches laid over the life of females for each combination of temperature and karyotype. (C) Longevity of both sexes in days for each combination of temperature and karyotype.

### Female fecundity is higher in *Cf-Inv(4*.*1)* NN homokaryotypes

Female fecundity varied significantly between inversion karyotypes, with an advantage of NN, regardless of temperature (Fig. 5A).

The number of eggs in the first clutch was mostly affected by the inversion karyotype (χ^2^ = 19.31, df = 1, p-value < 0.001). The NN females laid on average 144 ± 44 eggs in their first clutch compared to 97 ± 37 for SS females. The effect of the inversion interacted with continent (χ^2^ = 7.87, df = 1, p-value = 0.005). Indeed, the size of the first clutch for NN was significantly larger in North America (z = 5.26, p-value < 0.001) but not in Europe (z = 1.002, p-value = 0.75) although the same pattern was observed. Temperature did not have a significant impact on this trait (Fig. 5A, χ^2^ = 1.41, df = 2, p-value = 0.50). Mean clutch size followed the same trend, with significantly larger clutches for NN females without any interaction with temperature (see Supplementary Results 2, Supplementary Figure 8A).

The number of clutches laid by females during their life was not significantly impacted by the inversion karyotype (Fig. 5B, χ^2^ = 0.20, df = 1, p-value = 0.66). It was impacted by temperature (χ^2^ = 40.85, df = 2, p-value < 0.001), continent (χ^2^ = 16.63, df = 1, p-value < 0.001), and the interaction between the two (χ^2^ = 7.01, df = 2, p-value = 0.03), so that this global model explained 62.81% of the variance. North American females laid more clutches than European females (z = 3.54, p-value < 0.001), with an average of 2.83 ± 1.79 clutches in North America and 1.53 ± 1.52 in Europe. The number of clutches followed a thermal curve with an optimum at 20°C and a significant decrease at 30°C (Fig. 5B, z = 4.91, p-value < 0.001), and at 10°C (z = 4.48, p-value < 0.001). However, no significant effect of the interaction between inversion karyotype and temperature was detected (χ^2^ = 1.30, df = 2, p-value = 0.52, Fig 5B).

The total fecundity of a female is the product of the two traits previously analyzed: the number of clutches laid by the female and the size of these clutches. Therefore, total fecundity was significantly impacted by all the variables that had an effect on the number of clutches or the clutch size: temperature, karyotype, continent and their interactions (see Supplementary Results 3, Supplementary Figure 8B).

### Disentangling possible confounding factors

Besides the different genetic background across continents, which may explain part of the fitness variability between Europe and North America, we also controlled for the frequency of the chromosome I supergene, *Cf-Inv(1)*. For the North America continent, *Cf-Inv(1)* was unlikely to affect our results since its frequency did not significantly differ between North American NN and SS (Chi-squared test, χ^2^ = 0.02, df = 1, p-value = 0.878). However, the European experimental lineage SS displayed a considerably higher frequency of the arrangement α for *Cf-Inv(1)* than the NN lineage (Chi-squared test, χ^2^ = 26.54, df = 1, p-value < 0.001). Because the arrangement α of *Cf-inv (1)* reduces survival and increases fecundity (Mérot et al., 2020), it may have confounded some of the effects of *Cf-Inv(4)* identified in North America, and partially explain why no significant difference in fecundity was observed in Europe and why the reverse pattern was observed for viability.

## Discussion

Polymorphic inversions are increasingly being detected throughout the tree of life owing to the development of sequencing technologies, attracting a lot of attention in evolutionary biology due to their role in adaptation and diversification (Berdan et al., 2023). However, understanding the evolution of inversions requires not only genomic characterization but also knowledge about their distribution in time and space, as well as their functional impact on phenotypes (Lowry & Willis, 2010). Focusing on an inversion previously detected from genomic data in *C. frigida* (Mérot et al., 2021), our study provides indirect evidence that this inversion has a single evolutionary origin across North America and Europe where it shows parallel frequency clines along latitudinal eco-climatic gradients. Our experiments revealed an effect of *Cf-Inv(4*.*1)* on some life-history traits, partially interacting with temperature, but did not fully explain selection along the latitudinal cline. While the interaction between fitness, inversion, and temperature is complex, we suggest that the relative fitness advantage of the different karyotypes may vary with temperature, or other factors co-varying with latitude, and determine the local balance of inversion frequencies.

### A single origin for an inter-continental inversion polymorphism

Several lines of evidence show that the polymorphic inversion *Cf-Inv(4*.*1)*, previously described in North America (Mérot et al., 2021), is also polymorphic in Europe and shares a single origin across continents. First, genetic variation within the region of *Cf-Inv(4*.*1)* in Europe shows a high genetic structure in three clusters, confirming that the SNPs present in the region are differentiated and highly linked, as expected in the absence of recombination between the two arrangements. When measured with *F*_ST_, strong differentiation between the two arrangements is observed on both continents. In other words, the absence of recombination at the inversion level keeps the genetic divergence between the two arrangements far higher than the divergence caused by isolation by distance. In contrast, differentiation between continents is low within each arrangement of *Cf-Inv(4*.*1)*, with *F*_ST_ values closer to the rest of the genome. Overall, this leads to the clustering of the *Cf-Inv(4*.*1)* genomic region by arrangements instead of by continents, which supports a divergence of the two rearrangements before the split between North American and European populations, and suggests that the inversion *Cf-Inv(4*.*1)* has a single and old origin. While the age of the inversion is at least 61,000 – 134,000 years (Mérot et al., 2021), the geographic origin of the inversion and the history of continental colonizations remain unknown, and further biogeographic and demographic analysis would be needed to reconstruct past scenarios. Such a worldwide distribution is not unusual, with inversion polymorphisms being sometimes shared even across species boundaries (Knief et al., 2024; Tuttle et al., 2016). For example, Kapun et al. (2023) demonstrated that the inversion *In(3R)Payne* in *Drosophila melanogaster* originated in sub-Saharan tropical regions and subsequentially spread around the world, being maintained by selection on four continents. Several cosmopolitan inversions are also described in *Timema cristinae* stick-insects (Lindtke et al., 2017) and *Philomachus pugnax* ruffs (Küpper et al., 2016). In most cases, inversions that remain polymorphic for a long time and across all populations are likely to be under some form of balancing selection (Wellenreuther & Bernatchez, 2018). This balancing selection can result from different mechanisms including frequency-dependent selection, antagonistic pleiotropy, overdominance, disassortative mating and spatially/temporally variable selection (Cheng et al., 2012; Chouteau et al., 2017; Kim et al., 2017; Tuttle et al., 2016).

In *C. frigida*, the high divergence between arrangements led us to develop a marker with high operational concordance in North America (99%) but with a lower accuracy in Europe (84%). This imperfect concordance SNP-inversion is partly explained by shared polymorphism between European arrangements, possibly suggesting gene conversion events, which could have led to the transfer of a small track of DNA from one arrangement to the other one (Chen et al., 2007). For the remaining individuals, the inconsistencies are possibly due to protocol issues (amplification problems, incomplete digestion, etc). We acknowledge that this limitation somewhat reduces our confidence in the karyotyping of wild and experimental European individuals. In wild populations, this limitation was partially compensated by large sample sizes, while in experimental populations we cannot rule out that part of the different phenotypic patterns observed in Europe may be due to this incomplete association marker-inversion.

### Parallel clines of inversion frequencies along similar gradients

Because chromosomal inversions favor the long-term co-existence of ecotypes and can be under strong selection (Faria, Johannesson, et al., 2019), they are predicted to be frequently involved in repeated local adaptation across distant locations (Westram et al., 2022). For example, in the East African honeybees *(Apis mellifera)*, two genomic regions are highly divergent between highland and lowland populations of three distinct pairs, suggesting that these putative inversions are repeatedly implicated in adaptation to altitude (Christmas et al., 2019; Wallberg et al., 2017). This pattern can also be observed at larger geographical scale. The role of inversions in parallel adaptation is supported by theoretical simulations (Westram et al., 2022), as well as by numerous empirical studies across patchily distributed habitats like in the stick insect *Timema cristinae* and the Atlantic cod *Gadus morhua* (Lindtke et al., 2017; Matschiner et al., 2022), or along continuous environmental gradients as in sticklebacks and periwinkles (Jones et al., 2012; Morales et al., 2019).

Studying the frequencies of inversion *Cf-Inv(4*.*1)* in *C. frigida* at different locations along the European latitudinal gradient highlighted a cline that is parallel to the one identified on the Eastern coastline of North America by Mérot et al. (2021). On both continents, the latitudinal cline was in the same direction with the N arrangement more frequent in the North and the S arrangement more frequent in the South. This parallelism provides indirect evidence that the inversion is involved in adaptation to the same environmental gradient across continents (Bolnick et al., 2018; Westram et al., 2022).

Because several eco-climatic variables - such as photoperiod, temperature seasonality, desiccation, frequency of frost, but also often competition, diseases and parasitism prevalence (Cuevas et al., 2019) - co-vary with latitude, it is difficult to disentangle which factor may exert natural selection on the inversion. Temperature is a good candidate because it is one of the most stressful and influential abiotic factors for the survival all of living organisms, and especially for small ectotherms, such as insects (Denlinger & Lee Jr, 2010). Testing this hypothesis, we reported a significant relationship between inversion frequencies and annual mean air temperature on both continents. Such parallel association between inversion and temperature gradients is also observed in several other species, particularly in the genus *Drosophila*. For example, in *D. melanogaster*, four inversions form worldwide parallel latitudinal clines (Kapun & Flatt, 2019), suggesting that inversions may provide an important genetic architectural component for repeated adaptation along thermal gradients.

### Phenotypic impact of *Cf-Inv(4*.*1)* and temperature

Although the implication of inversions in adaptation to different environments is increasingly suspected or demonstrated in a large number of species, pinpointing the underlying traits and functional mechanisms remains challenging and requires a direct, experimental approach. Adaptation along a climatic gradient may in fact involve a wide range of traits. Insects are particularly sensitive to temperature, which affects their survival at all life stages and fecundity (Régnière et al., 2012). Typically, each species has an optimal range of temperature and higher temperature induces lower survival while lower temperature delays development (Hoffmann et al., 2003). Although thermal tolerance, often assessed through critical thermal limits, has traditionally been used to predict climate adaptation, and species persistence with climate change, recent studies suggest that reproductive thermal limits may provide a more accurate predictor of vulnerability. These limits could offer a finer measure of species thermal susceptibility than survival (Parratt et al., 2021; van Heerwaarden & Sgrò, 2021). Our results in *C. frigida* confirm that temperature plays an important role in shaping several life-history traits. The life cycle duration of *C*.

*frigida* increased when temperature decreased, with a longer development time but also a longer longevity. Viability and fecundity (total fecundity and number of clutches) were also strongly affected by temperature, forming classical bell-shaped thermal performance curves. At the thermal limits, insects may display different responses including reversible loss of motor function (hot/cold coma). Interestingly, some of those traits are associated with inversions in other species, adding support for a causal link between these traits and climatic gradients. For example, in the melon thrips *Thrips palmi*, Ma et al. (2024) experimentally demonstrated the role of three inversions in range expansion across a climate gradient mediated by an effect on the maximal critical temperature (CTmax). In the mosquito *Anopheles gambiae*, the chromosomal inversion 2La, which form aridity clines in Africa, is associated with increased survival to heat stress following heat hardening (Rocca et al., 2009).

In *C. frigida*, we did not observe any effect of *Cf-Inv(4*.*1)* on cold tolerance, with our selected metrics, but we did report, for all karyotypes, significant tolerance to prolonged exposure to subzero temperatures (16h at -5°C) and deep supercooling ability (ranging from -7.8°C to -23.9°C). This resistance suggests that *C. frigida* follows a freeze-avoidance strategy, specifically classified as “chill-tolerant” according to Bale’s classification (Bale, 1993). Chill-tolerant species can endure extended exposure to sub-zero temperatures by remaining in a supercooled state, but they risk mortality when temperatures exceed their supercooling point (SCP). In those species, survival and body condition is thus more affected by prolonged chilling rather than by extreme freezing events (Andersen et al., 2015). To fully characterize thermal tolerance in *C. frigida*, a more comprehensive approach is likely needed, such as the Thermal Death Time (TDT) methods, which combines many intensities of stress at many durations (Jørgensen et al., 2022; Rezende et al., 2014).

*Cf-Inv(4*.*1)* exhibits significant impacts on several life-history traits, in interaction with thermal conditions. Although these observed patterns differed between continents, the overall results combining the whole dataset indicate that the *Cf-Inv(4*.*1)* seems to be involved in a life-history trade-off impacting both viability and fecundity. Egg-to-adult viability varied with temperature and inversion karyotypes, forming classical performance curves, with overall broader amplitude of tolerance and higher survival in SS karyotypes compared to NN karyotypes. Female fecundity, however, was higher for NN than SS, suggesting that the inversion *Cf-Inv(4*.*1)* may strongly influence fitness.

Parallelism across continents in the inversion-phenotype relationship can, nevertheless, not be confirmed by our dataset. For fecundity, the signal is in the same direction on both continents but not significant in Europe while viability shows a European pattern opposite to that observed in North America. On top of a different genetic background between the two continents, the inversion *Cf-Inv(1)*, which is known to impact fecundity and viability (Mérot et al., 2020), may have acted as a confounding factor in European experimental populations. In fact, although a complete karyotyping of all individuals would have been more accurate, our estimates based on a subset of samples suggest that the inversion *Cf-Inv(1)* was at different frequencies between Europe-SS and Europe-NN lines (but at similar frequencies between North America-SS and North America-NN). We thus cannot rule out that the phenotypic effect of *Cf-Inv(1)* contributed to some of the differences observed between NN and SS in Europe. Since our experiments suggest that the two inversions impact similar traits (viability, fecundity), further experiments would be required to fully understand putative additive and epistatic interactions, although it is worth noting that the two inversions segregate independently in wild populations (linkage disequilibrium = 0.03 (North America) & 0.02 (Europe)).

Overall, the inversion *Cf-Inv(4*.*1)* tends to have opposing effects on different fitness traits, possibly placing the two karyotypes at different levels in the trade-off between surviving and reproducing. This property is shared by several large inversion supergenes, possibly due to the high number of genes linked together or to the accumulation of deleterious mutations, and it is supported by numerous examples in the literature, such as seen in zebra finches (Pei et al., 2023), in *Heliconius numata* butterflies (Jay & Joron, 2022) and in stick insects (Nosil et al., 2023). Indeed, if the inversion links an additive beneficial allele with a recessive deleterious allele, polymorphisms can be maintained through antagonistic pleiotropy of the entire inversion haplotype (Pei, 2021). This mechanism often translates into higher fitness in heterozygotes, or overdominance, as is the case in stick insects and zebra finches. Further experiments to measure fitness of heterokaryotypes in *C. frigida* could therefore help to deepen our understanding of the factors that contribute to the maintenance of the inversion *Cf-Inv(4*.*1)* polymorphism.

### The challenge of understanding the clinal pattern in the light of functional impacts

While the parallel clines in inversion frequency support a role for spatially variable selection along the eco-climatic gradients, laboratory experiments highlight antagonistic fitness effects for broad conditions of temperature. This may reflect the fact that inversion evolution is driven by multiple interacting processes occurring at the same time because large inversions encompass hundreds of genes and impact a broad range of phenotypes (Berdan et al., 2023). For instance, Nosil et al. (2023) demonstrated that an inversion polymorphism in *Timema* stick insects is maintained by a combination of processes, including life-history trade-offs, heterozygote advantage, local adaptation to different hosts, and gene flow.

The cline of inversion frequencies observed in the wild led us to expect the southern arrangement to be favored under high temperatures and the northern arrangement to be favored under low temperatures. Although experiments revealed an effect of *Cf-Inv(4*.*1)* on some life-history traits, we did not detect sufficient interaction with temperature to explain the cline. It is worth noting that beside constant exposures, the effect of temperature could act through diverse variables such as temperature seasonality, daily fluctuations, or even in combination with other climatic variables (precipitations, season length, day length). For instance, Xing & Zhao (2022) demonstrated the effect of daily temperature amplitudes on longevity, fecundity and the thermal tolerance in adult diamondback moths. Furthermore, the cline could be explained by inversion-latitude effects on fitness traits that were not measured, like mating or foraging activity under varying conditions of light and temperature (Rivas et al., 2018). Indeed, exploring the genomic surroundings of *Cf-Inv(4*.*1)* has identified tentative candidate genes possibly underlying phenotypic variation along a latitudinal gradient in circadian rhythm that appears to interact with temperature (Fig. 2G, see also genome annotation in Mérot et al., 2021). Among the candidates, we found *regucalcin*, a gene paralogous to the *Drosophila spp*. cold acclimatation gene (*Dca*), which is involved in the adaptive response to low temperatures (Arboleda-Bustos & Segarra, 2011). We also found *bric à brac*, whose expression level varies with temperature in *Drosophila melanogaster* (De Castro et al., 2018), and which is a key regulator of sex pheromone choice in European corn borer moth males, strongly influencing mating behaviors (Unbehend et al., 2021). It is also worth noting the proximity of additional genes involved in the circadian rhythm, such as *period* and *tilB*, which also mediate the interaction between temperature and the biological clock (Kaiser et al., 2016; Sehadova et al., 2009), with *period* showing a latitudinal cline in *Drosophila* (Sawyer et al., 1997). *Cf-Inv(4*.*1)* further contains some genes known to be involved in fecundity, such as *Asterix* (Ipsaro & Joshua-Tor, 2022) and *Notch* (Xie et al., 2024), which could explain the effects on clutch size in our experiments. Xie et al. (2024) even demonstrated experimentally that *Notch* affects mating frequency and egg production in *Basilepta melanopus*. Future deeper understanding of the functional effects of allelic variation associated with each arrangement, as well as their conservatism or novelty across continents, could be inferred by further investigating the putative candidate genes or quantifying differential gene expression (*e*.*g*. (Fuller et al., 2016; Said et al., 2018) between arrangements and under different climatic conditions.

Classically, frequency clines are thought to result from an equilibrium between selection and migration. However, Barton & Gale (1993) mentioned an alternative model, according to which a cline can be formed by different mechanisms of balancing selection if the equilibrium of genotype proportions varies according to the environment. According to Westram et al. (2022), this scenario is particularly relevant for stable, large-scale clines, as it allows for the combination of strong selection with subtle frequency gradients. For instance, this model is strongly suspected to underlie parallel clines in inversion *In(3R)Payne* in *D. melanogaster*, which display gradual frequency changes across a broad geographic range (Kapun & Flatt, 2019; Westram et al., 2022). We therefore speculate that, in the context of a trade-off between fecundity (higher in NN) and thermal stress tolerance (higher in SS), the relative fitness value of both arrangements of *Cf-Inv(4*.*1)* may differ along the latitudinal thermal gradient, so that frequencies at equilibrium vary clinally. It is possible that high latitude conditions, such as low temperature but also shorter days or a shorter season, select for a higher investment in fecundity which would increase fitness value of arrangement N. Conversely, environmental conditions at low latitude, such as higher temperature or stronger fluctuations of temperature due to a thinner and less-buffered wrack bed habitat, in combination with competition and desiccation stress, could select for an enhanced survival during larval phase, which would increase the fitness value of arrangement S. Overall, such as scenario of a balanced cline (Barton & Gale, 1993) is more plausible for a polymorphic cline, where neither allele reaches fixation at either end of the cline, as we observed for *Cf-Inv(4*.*1)*. As this kind of inversion under balancing selection is expected to be polymorphic over a wide range, it is also more probable that the colonization of a new environmental gradient, even by a small founding group, results in the appearance of a parallel large-scale cline (Westram et al., 2022). To further address the role of chromosomal inversions in parallel adaptive processes, it will be important to trace the evolutionary and demographic history of rearrangements and to understand intra-inversion diversity (*cf*. (Kim et al., 2022; Knief et al., 2024; Lundberg et al., 2023).

## Supporting information

Metadata ENA Samples

Metadata NCBI Samples

Data Clines

Data Development Viability

Data Fecundity

Data Longevity

Script Cline Analyses

Script Fitness Analyses

## Data Availability

Data supporting this study are available in supplementary materials.

Whole genome sequences of European individuals can be found on ENA (PRJEB86057). A data frame with metadata for all individuals including SRA accession numbers is available in supplementary materials (file Metadata_ENA_samples.csv).

Whole genome sequences of North American individuals can be found on NCBI (PRJNA689963). A data frame with metadata for all individuals including SRA accession numbers is available in supplementary materials (file Metadata_NCBI_samples.csv).

The reference genome assembly is available on NCBI (PRJNA688905).

Scripts used to run the genomic analyses are available at https://github.com/clairemerot/angsd_pipeline. Scripts used to run the statistical analyses on experimental data is available in supplementary materials.

## Acknowledgements

We thank Emmanuel Le Rouzic and Astrid Bavay for help with the experiments and the platform ECOLEX (Ecobio), Manon Norest, Christine Peré and Charles Babin for help with molecular biology and the platforms PEM (Ecobio) and PAG (Université Laval). We thank Nauras Daraghmeh for his inputs in the project and data management. We are very grateful to Louis Bernatchez for earlier inputs in the project and support for sequencing.

## Author Contribution Statements

LN: Conceptualization, Formal analyses, Investigation, Project administration, Visualization, Writing – original draft.

EB: Conceptualization, Funding acquisition, Reviewing the draft.

MW : Conceptualization, Funding acquisition, Reviewing the draft.

PDW: Conceptualization, Funding acquisition, Reviewing the draft.

HC: Conceptualization, co-supervision, Reviewing the draft.

AC: Formal analyses, Investigation, Visualization.

SG: Conceptualization, co-supervision, Reviewing the draft

CM: Conceptualization, Funding acquisition, Investigation, Methodology, Project administration, Supervision, Writing – original draft.

