## Supplementary Material for "Parallel clines of chromosomal inversion frequencies in seaweed flies are associated with thermal variation"

#### Summary

Supplementary methods 4. PCR and Digestion protocol for genotyping with diagnostic marker . 2
Supplementary figure 3. Genotypes at all biallelic SNPs within inversion *Cf-Inv(4.1)* between positions 1,634,841 (3 kb before diagnostic SNPs) and 1,640,843 (3 kb after diagnostic SNPs).. 10
Supplementary figure 4. Clines of inversion frequency with whole-genome sequencing data. .. 11

### Supplementary Methods

#### Supplementary methods 1. Laboratory breeding of flies

Flies were raised in plastic boxes of 1.7L (13 × 8.5 × 18 cm) containing approximately 50% of Fucacea (Fucus serratus and/or Fucus vesiculous) and 50% of Laminariaceae (Saccharina latissima and/or Laminaria digitata). At each generation, emerging adults were captured over a few days using an insect vacuum and presented with fresh seaweed to stimulate egg-laying. A random subsample of eggs was then collected (to control for density) and spread on the same natural seaweed substrate in two to three boxes per population. Each population was started with 100-300 individuals to ensure enough genetic diversity, and each generation usually produced 100-300 adults. The laboratory populations were kept for 7 to 14 generations before starting the experiments.

#### Supplementary methods 2. Extraction protocol for European WGS data

To perform a PCA and to validate the use of the diagnostic marker in Europe, we used the DNA of 56 Scandinavian individuals extracted by Emma Berdan in 2015.

DNA was extracted using a modified version of the PureGene (Gentra Systems, www.gentra.com) extraction protocol over four days. On the first day, tissue samples were placed in 600 µl of cell lysis solution (0.1 M Tris, 0.0077 M EDTA, and 0.0035 M sds) with 3 µl of Proteincase K (20 mg/ml). The sample was vortexed and kept at 65° overnight. On the second day, 200 µl of protein precipitation solution (Qiagen, Valencia, CA) was added and the sample was vortexed and then stored at 4° C overnight. On day three, the sample was centrifuged at 12.6 x $10^3$ rpm for 5 minutes and then the supernatant was removed leaving behind the protein pellet. Six hundred µl of isopropanol was added and the sample was kept at -20° overnight. On the final day, the sample was centrifuged at 12.6 x $10^3$ rpm for 4 minutes to precipitate DNA. The supernatant was removes and 600 µl of 70% ethanol was added. The sample was vortexed and then centrifuged again. The ethanol was removed and the pellet was allowed to dry and then rehydrated with 30 %l of TE. Sample concentration and quality were verified using a Nanodrop spectrophotometer.

#### Supplementary methods 3. Quicklysis extraction protocol

DNA extraction for genotyping was realized as follows. The flies' heads were delicately detached from the rest of the body and let dried to get rid of ethanol. Each head was then placed for digestion in a solution composed of 1µL of proteinase K (20mg/mL) and 50µL of a lysis buffer made of 0.5% Tween 20, 0.04M of Tris-HCl (pH 9.0), and 0.05M of KCl. The lysis mix and the head were incubated 12 hours at 37°C to allow the digestion of the cells and 15 minutes at 95°C in a thermocycler to inactivate the proteinase. Once the incubation completed, the extraction product was centrifuged during 12 minutes at 4000rpm in order to obtain a clear lysate, and then used immediately or stored at -20°C until PCR.

#### Supplementary methods 4. PCR and Digestion protocol for genotyping with diagnostic marker

The PCR was performed in a 25µL final volume, containing 14.75µL of sterile H2O, 5µL of a taq buffer, 1µL of the forward primer F634 whose sequence is AGAATCTCCGTGCCATGCAA, 1µL of the reverse primer R1301 whose sequence is GCACCTTGCAAGCCATCTTC, 0.25µL of taq polymerase (Meridian society) and 3µL of the supernatant obtained after centrifugation of the extraction product. The PCR program was as follows: 2 minutes at 95°C for the initial denaturation; 35 cycles consisting of 45 seconds at 95°C, 45 seconds at 55°C, and 1 minute at 72°C; and finally, 15 minutes at 72°C for the final elongation. The PCR product was then stored at 4°C until digestion. Only 103 out of the 104 North American samples and 38 individuals out of 56 European samples had enough DNA left to perform a successful PCR. For the digestion by EcoRI, 5µL of PCR product were added to 10µL of a solution containing 0.5µL of the enzyme (20000U/mL, New England BioLabs), 2.5µL of Cutsmart buffer 10X (New England BioLabs) and 7µL of sterile water. The digestion program was set to 30 minutes at 37°C. The digestion product was then run on 2.5% agarose gel electrophoresis for 45 minutes at 115V (figure X).

##### Supplementary methods 5. Measuring supercooling point

To measure the supercooling point (SCP), we exposed flies to decreasing temperature in a cold oil bath and measured temperature with a thermocouple, *i.e.*, a very thin sensor that records the temperature at regular intervals. Adult flies were maintained alive in contact with a thermocouple type K with parafilm. The thermocouple equipped with a fly was placed at the bottom of a glass tube prevented from moving with a piece of cotton. The thermocouple was connected to a Testo 176T4 temperature data logger (Testo SE& Co., Germany). The tube was then immersed in the mineral oil bath of a cryostat bath (Polystat CC3, Huber Kältemaschinenbau AG, Germany) previously set to 5°C. A one-hour program was launched to gradually lower the temperature to -25°C at a decreasing rate of 0.5°C/min. The temperature of the insects was recorded every second. The SCP was defined as the temperature at the onset of the freezing exotherm produced by the latent heat. Once the program was complete, the frozen/dead flies were gently detached to be sexed and weighed. Adult flies were stored in 96% ethanol until genetic analysis. Our experimental design allows to test 8 flies at the same time. Flies were chosen randomly among freshly-emerged adults (1-2 days old) kept in semi-natural conditions. We tested 56 to 64 flies per location for a total of 288 flies.

##### Supplementary methods 6. Measuring chill coma recovery time

To measure Chill Coma Recovery Time (CCRT), 20 flies from the same location were placed in a glass tube, immersed in the oil bath of the cryostat at - 5°C. After 16 hours the flies were removed from the tubes and placed on their backs in a room at 15°C. This temperature × duration was chosen based on a series of preliminary assays which showed that a temperature of – 10°C was too low and causes the freezing (and death) of some flies and that a duration of few hours was too short to observe sufficient variability between individuals. The CCRT was determined as the time between the flies exiting the cold bath and the moment they turned back on their legs. To account for the duration of this turn and allow accurate flies sorting, these times were classified in 30-second interval classes. The flies were then weighed, sexed, and stored in 96% ethanol until genotyping.

#### Supplementary Results

##### Supplementary results 1. Cold tolerance

Contrary to our predictions, the SCP was affected neither by the location ($F = 1.79$, $df = 4$, p-value = 0.13), nor by the inversion *Cf-Inv(4.1)* ($F = 0.98$, $df = 2$, p-value = 0.38). SCP was significantly associated with mass, with larger flies freezing at lower temperatures ($F = 5.80$, $df = 1$, p-value = 0.017). We also detected a significant effect of the inversion *Cf-Inv(1)* in interaction with sex ($F = 6.22$, $df = 2$, p-value = 0.002), ββ males having a slightly lower

SCP than other combinations. However, those effects were marginal and the model explained only 6.3% of the high variance in SCP (min = -23.9°C, max = -7.8°C, sd = 3.48°C).

The CCRT was significantly impacted by the origin location of the flies ($\chi^2 = 16.49$, df = 4, p-value = 0.002), but counter-intuitively, flies from a southern location (MA) recovered slightly faster than flies from RB (Chi-squared = 7.3, df = 1, p=0.007), PT (Chi-squared = 9.7, df = 1, p=0.002) and NH (Chi-squared = 5.2, df = 1, p=0.02) locations. We also detected a significant effect of inversion *Cf-Inv(1)* on CCRT ($\chi^2 = 8.63$, df = 2, p = 0.013), with ββ flies recovering slightly slower than αα flies (log-rank test, $\chi^2 = 4.1$, df = 1, p-value = 0.04) and αβ flies (log-rank test, $\chi^2 = 7.5$, df = 1, p-value = 0.006). The model explained 18.7% of the variance in CCRT.

#### Supplementary results 2. Mean clutch size

Mean clutch size was mostly impacted by the inversion karyotype ($\chi^2 = 15.01$, df = 1, p-value < 0.001). The effect of this variable was also exerted in interaction with the continent ($\chi^2 = 5.52$, Df = 1, p-value = 0.019). Indeed, NN females laid on average significantly larger clutches than SS in North America (z = 4.71, p-value < 0.001) but not in Europe (z =1.13, p-value = 0.67). However, no interaction between inversion and temperature was detected ($\chi^2 = 4.06$, df = 2, p-value = 0.13).

#### Supplementary results 3. Total fecundity

The total fecundity of a female is the product of two other traits: the number of clutches laid by the female and the size of these clutches. Therefore, analyses performed on total fecundity identified significant effects of all the variables that had an effect on the number of clutches or the clutch size. Indeed, Temperature ($\chi^2 = 49.88$, Df = 2, p-value < 0.001), continent ($\chi^2 = 33.73$, Df = 1, p-value < 0.001) and inversion karyotype ($\chi^2 = 7.63$, Df = 1, p-value = 0.0058) were all significant with considerable effects on the total number of eggs laid by females. The total fecundity was higher at 20°C than at 30°C (z = 5.39, p-value < 0.001) and 10°C (z = 4.35, p-value < 0.001) and was higher in North America than in Europe (z = 4.36, p-value < 0.001). We also observed that females of the NN karyotype for the inversion laid more eggs in their life than SS females. The total contribution of the model reaches 64% of explained variance when taking into account the interactions between factors. Indeed, when separating the data according to the continent, we observed that this effect of karyotype was significant in North America (z = 3.11, p-value = 0.01) but not in Europe (z = 0.087, p-value = 0.99). However, we couldn't detect any effect of interaction between inversion and temperature ($\chi^2 = 1.8$, df = 2, p-value = 0.41).

#### Supplementary results 4. Analyses of marker inconsistencies

In North America, the diagnostic marker showed a very good accuracy with 99% concordance with inversion karyotypes determined by PCA (Mérot et al., 2021). Out of 103 individuals genotyped with the marker, 102 were consistent with the karyotype obtained from the PCA on whole genome sequences. One individual genotyped as SS by the PCA was assigned as SN by the marker (Supplementary Table 3). In Europe, 32 out of 38 individuals were consistent, representing a concordance rate of 84%. Most of the remaining individuals (5) were assigned as heterokaryotes by the PCA but as homokaryotes by the diagnostic marker (4 SS and 1 NN). One individual was also assigned NN by the PCA and heterokaryote by the diagnostic marker (Supplementary Figure 3).

The discordance could be explained either by the fact that a high number of missing data in whole genome sequences tends to place an individual among heterokaryotypes on the PCA, or by a gene conversion event, which could have led to the transfer of part of one arrangement to the other one, resulting in inconsistencies between the genotyping of this specific region and the global pattern of the entire inversion (Chen et al., 2007). Although the sliding-window PCA did not highlight any significant shifts in the PC1 score of the individuals concerned by the inconsistencies, it is important to note that gene conversion events are generally shorter than 1kb in D. melanogaster (Miller et al., 2012) and are unlikely to be detected in 50kb windows.

To examine the cause of the diagnostic marker inconsistencies, we plotted the genotypes of all SNPs within a region of 6 kb around the two diagnostic SNPs (see also Supplementary Figure 3). This analysis revealed that in the six incorrectly genotyped individuals in Europe, four carry a small segment around the diagnostic SNPs, in which the genotypes of the SNPs are inconsistent with the rest of the inversion. Three of them harbor the same segment of 189 bp with SNP genotypes corresponding to the S arrangement on an N haplotype. One of them harbors a segment of 508 bp with SNP genotypes corresponding to the N arrangement on a S haplotype. This additional polymorphism is consistent with patterns of gene conversion. For the remaining individuals, the inconsistencies are possibly due to protocol issues (amplification problems, incomplete digestion, etc).

#### Supplementary Tables

##### Supplementary table 1. North American sampling locations, number of flies used for supercooling point (SCP) and Chill coma recovery time (CCRT) experiments

| Location | Latitude | Longitude | N for SCP | N for CCRT |
|---|---|---|---|---|
| RB | 50.28161 | -65.51516 | 56 | 58 |
| PT | 49.31839 | -67.38624 | 56 | 59 |
| NB | 46.46795 | -62.41561 | 56 | 58 |
| MA | 41.92654 | -70.54451 | 64 | 76 |
| NH | 42.92081 | -70.79809 | 56 | 60 |
| CB | 45.12965 | -63.44673 | 0 | 0 |

##### Supplementary table 2. Sequences of primers used in the diagnostic assay

| | |
|---|---|
| F634 | AGAATCTCCGTGCCATGCAA |
| R1301 | GCACCTTGCAAGCCATCTTC |

#### Supplementary table 3. Whole-genome PCA versus diagnostic-marker karyotypes.

Column 1 and 2 contain the locations and continents of origin of the flies. Column 3 contain the karyotypes determined from PCA on whole-genome sequencing data in the region of *Cf-Inv(4.1)*. The column 4 contain the karyotypes determined after genotyping with the diagnostic marker.

| Location | Cont | PCA | Marker |
|---|---|---|---|
| Justoya | Eur | NN | SN |
| Justoya | Eur | SN | Non amplified |
| Justoya | Eur | NN | Unreadable |
| Justoya | Eur | SN | SN |
| Justoya | Eur | SN | SN |
| Justoya | Eur | NN | Non amplified |
| Justoya | Eur | SN | SN |
| Justoya | Eur | NN | NN |
| Justoya | Eur | SN | SN |
| Justoya | Eur | SN | Unreadable |
| Magnarp | Eur | SN | SS |
| Magnarp | Eur | SS | SS |
| Magnarp | Eur | SN | SN |
| Magnarp | Eur | SS | Non amplified |
| Magnarp | Eur | SN | Non amplified |
| Magnarp | Eur | SS | SS |
| Magnarp | Eur | SN | SN |
| Magnarp | Eur | NN | NN |
| Magnarp | Eur | SS | SS |
| Magnarp | Eur | SN | SN |
| Oldberg | Eur | SN | SN |
| Oldberg | Eur | NN | Non amplified |
| Oldberg | Eur | SN | Non amplified |
| Oldberg | Eur | SN | NN |
| Oldberg | Eur | SN | Non amplified |
| Oldberg | Eur | SN | Non amplified |
| Skeie | Eur | NN | Non amplified |
| Skeie | Eur | SN | Non amplified |
| Skeie | Eur | SN | Non amplified |
| Skeie | Eur | NN | NN |
| Skeie | Eur | NN | NN |
| Skeie | Eur | NN | Non amplified |
| Skeie | Eur | SN | Non amplified |
| Skeie | Eur | NN | NN |
| Skeie | Eur | NN | Non amplified |
| Skeie | Eur | SS | SS |
| Smygehuk | Eur | SS | Non amplified |
| Smygehuk | Eur | NN | NN |
| Smygehuk | Eur | SN | SN |
| Smygehuk | Eur | SN | SN |
| Smygehuk | Eur | SN | SN |
| Smygehuk | Eur | SN | SS |
| Smygehuk | Eur | SN | SN |
| Smygehuk | Eur | SS | SS |
| Smygehuk | Eur | SS | SS |
| Smygehuk | Eur | SN | SS |
| Stavder | Eur | SS | SS |
| Stavder | Eur | SN | SS |
| Stavder | Eur | SS | SS |
| Stavder | Eur | SN | SN |
| Stavder | Eur | SN | SN |
| Stavder | Eur | SN | SN |
| Stavder | Eur | SN | SN |
| Stavder | Eur | SN | SN |
| Stavder | Eur | SN | Unreadable |
| Stavder | Eur | SS | SS |
| AG | Ame | NN | NN |
| AG | Ame | SN | SN |
| AG | Ame | SS | SS |
| AG | Ame | SS | SS |
| AG | Ame | NN | NN |
| AG | Ame | SN | SN |
| AG | Ame | NN | NN |
| AG | Ame | SS | SS |
| AG | Ame | SN | SN |
| AG | Ame | NN | NN |
| AG | Ame | SS | SS |
| AG | Ame | SN | SN |
| BS | Ame | NN | NN |
| BS | Ame | NN | NN |
| BS | Ame | SN | SN |
| BS | Ame | SS | SS |
| BS | Ame | SN | SN |
| BS | Ame | SN | SN |
| BS | Ame | SN | SN |
| BS | Ame | NN | NN |
| BS | Ame | NN | NN |
| BT | Ame | SN | SN |
| BT | Ame | NN | NN |
| BT | Ame | SN | SN |
| BT | Ame | SN | SN |
| BT | Ame | NN | NN |
| BT | Ame | SS | SS |
| BT | Ame | NN | NN |
| BT | Ame | NN | NN |
| BT | Ame | SS | SS |
| BT | Ame | SS | SS |
| BT | Ame | SN | SN |
| BT | Ame | SS | SN |
| CB | Ame | NN | NN |
| CB | Ame | SN | SN |
| CB | Ame | SS | SS |
| CB | Ame | SS | SS |
| CB | Ame | NN | NN |
| CB | Ame | SS | SS |
| CB | Ame | SS | SS |
| CB | Ame | SN | SN |
| CB | Ame | SN | SN |
| CB | Ame | NN | NN |
| CB | Ame | SN | SN |
| CB | Ame | NN | NN |
| HA | Ame | NN | NN |
| HA | Ame | NN | NN |
| MA | Ame | SS | SS |
| MA | Ame | SS | SS |
| MA | Ame | SN | SN |
| MA | Ame | SS | SS |
| MA | Ame | SS | SS |
| MA | Ame | NN | NN |
| MA | Ame | NN | NN |
| MA | Ame | SN | SN |
| MA | Ame | SN | SN |
| MA | Ame | SN | SN |
| NB | Ame | SS | SS |
| NB | Ame | SN | SN |
| NB | Ame | SS | SS |
| NB | Ame | NN | NN |
| NB | Ame | SN | SN |
| NB | Ame | NN | NN |
| NB | Ame | SS | SS |
| NB | Ame | NN | Non amplified |
| NB | Ame | SN | SN |
| NB | Ame | SS | SS |
| NB | Ame | NN | NN |
| NB | Ame | SN | SN |
| RB | Ame | SS | SS |
| RB | Ame | NN | NN |
| RB | Ame | NN | NN |
| RB | Ame | SS | SS |
| RB | Ame | SN | SN |
| RB | Ame | SN | SN |
| RB | Ame | NN | NN |
| RB | Ame | SN | SN |
| RB | Ame | SS | SS |
| RB | Ame | SN | SN |
| RB | Ame | NN | NN |
| RC | Ame | SN | SN |
| RC | Ame | NN | NN |
| RC | Ame | SN | SN |
| RC | Ame | SS | SS |
| RC | Ame | SN | SN |
| RC | Ame | SN | SN |
| RC | Ame | NN | NN |
| RC | Ame | SS | SS |
| RC | Ame | NN | NN |
| RC | Ame | NN | NN |
| RC | Ame | SS | SS |
| RC | Ame | SS | SS |
| SI | Ame | SS | SS |
| SI | Ame | SN | SN |
| SI | Ame | SN | SN |
| SI | Ame | SS | SS |
| SI | Ame | SN | SN |
| SI | Ame | NN | NN |
| SI | Ame | NN | NN |
| SI | Ame | NN | NN |
| SI | Ame | SS | SS |
| SI | Ame | NN | NN |
| SI | Ame | SS | SS |
| SI | Ame | SN | SN |

#### Supplementary table 4. Estimated parameters of thermal performance curves for viability.

Rmax is the estimated maximal viability. Topt is the estimated optimal temperature. Ctmin is the estimated minimal critical temperature. Ctmax is the estimated maximal critical temperature. Thermal tolerance is the amplitude of temperatures for which performance is above zero. Breadth is the amplitude of temperatures for which performance is above 80% of the rmax. A skewness of zero means that the thermal performance curve is symmetrical. Group is the combination of karyotype (SS/NN) and continent (A = North America / E = Europe).

| rmax | topt | ctmin | ctmax | e | eh | q10 | thermal safety margin | thermal tolerance | breadth | skewness | group |
|---|---|---|---|---|---|---|---|---|---|---|---|
| 0,34 | 22,29 | 9,16 | 35,43 | 0,9 | -0,25 | 3,46 | 13,14 | 26,27 | 11,76 | 1,15 | SS_A |
| 0,16 | 21,49 | 9,97 | 33,01 | 1,56 | 0,66 | 8,61 | 11,52 | 23,04 | 10,34 | 0,9 | NN_A |
| 0,18 | 20,41 | 9,42 | 31,4 | 0,84 | 1,08 | 3,18 | 10,99 | 21,98 | 9,86 | -0,25 | SS_E |
| 0,3 | 21,56 | 10,14 | 32,98 | 1,35 | 1,06 | 6,49 | 11,42 | 22,84 | 10,24 | 0,3 | NN_E |

Supplementary table 5. Summary of Generalized Linear Models

| Model N° | 1 | | | 2 | | | 3 | | | 4 | | | 5 | | | 6 | | |
|---|---|---|---|---|---|---|---|---|---|---|---|---|---|---|---|---|---|---|
| Response variable | Viability | | | Development time | | | Development time 15°C | | | Development time 20°C | | | Development time 25°C | | | Development time 30°C | | |
| Type | glm | | | glm | | | glm | | | glm | | | glm | | | glm | | |
| Distribution | Binomial (link = logit) | | | Gamma | | | Gamma | | | Gamma | | | Gamma | | | Gamma | | |
| | Chi² | Df | p-value | Chi² | Df | p-value | Chi² | Df | p-value | Chi² | Df | p-value | Chi² | Df | p-value | Chi² | Df | p-value |
| Karyotype | 12,6 | 1 | *** | 0 | 1 | NS | 0,38 | 1 | NS | 2,57 | 1 | NS | 4,59 | 1 | * | 0,87 | 1 | NS |
| Temperature | 68,48 | 3 | *** | 49566 | 3 | *** | | | | | | | | | | | | |
| Continent | 14,07 | 1 | *** | 0 | 1 | NS | 0,24 | 1 | NS | 1,39 | 1 | NS | 0,5 | 1 | NS | 0,006 | 1 | NS |
| Sexe | | | | 414 | 1 | *** | 161,85 | 1 | *** | 250,69 | 1 | *** | 102,13 | 1 | *** | 97,32 | 1 | *** |
| Karyotype : Temperature | 35,27 | 3 | *** | 10 | 3 | * | | | | | | | | | | | | |
| Karyotype : Continent | 174,05 | 1 | *** | 13 | 1 | *** | 10,65 | 1 | ** | 2,93 | 1 | NS | 2,59 | 1 | NS | 0,61 | 1 | NS |
| Temperature : Continent | 10,24 | 3 | * | 0 | 3 | NS | | | | | | | | | | | | |
| Karyotype : Sexe | | | | 6 | 1 | * | 11,01 | 1 | *** | 0,01 | 1 | NS | 0,94 | 1 | NS | 0,001 | 1 | NS |
| Temperature : Sexe | | | | 173 | 3 | *** | | | | | | | | | | | | |
| Continent : Sexe | | | | 0 | 1 | NS | 0,09 | 1 | NS | 0,14 | 1 | NS | 0 | 1 | NS | 2,39 | 1 | NS |
| Classic R² | 0,43 | | | 0,98 | | | 0,36 | | | 0,41 | | | 0,32 | | | 0,33 | | |
| Deviance R² | 0,43 | | | 0,98 | | | 0,36 | | | 0,4 | | | 0,34 | | | 0,33 | | |

| Model N° | 7 | | | 8 | | | 9 | | | 10 | | | 11 | | |
|---|---|---|---|---|---|---|---|---|---|---|---|---|---|---|---|
| Response variable | First clutch size | | | Mean clutch size | | | Number of clutches | | | Total fecundity | | | Longevity | | |
| Type | glm | | | glm | | | glm | | | glm | | | glm | | |
| Distribution | Quasi (link = identity) | | | Quasi (link = identity) | | | Poisson | | | Quasi-Poisson | | | Gamma | | |
| | Chi² | Df | p-value | Chi² | Df | p-value | Chi² | Df | p-value | Chi² | Df | p-value | Chi² | Df | p-value |
| Karyotype | 19,31 | 1 | *** | 15,01 | 1 | *** | 0,2 | 1 | NS | 7,63 | 1 | ** | 0,7 | 1 | NS |
| Temperature | 1,41 | 2 | NS | 3,34 | 2 | NS | 40,85 | 2 | *** | 49,88 | 2 | *** | 798,9 | 2 | *** |
| Continent | 3,61 | 1 | NS | 1,49 | 1 | NS | 16,66 | 1 | *** | 33,73 | 1 | *** | 16,46 | 1 | *** |
| Sexe | | | | | | | | | | | | | 8,06 | 1 | ** |
| Karyotype : Temperature | 2,28 | 2 | NS | 4,06 | 2 | NS | 1,3 | 2 | NS | 1,8 | 2 | NS | 1,21 | 2 | NS |
| Karyotype : Continent | 7,87 | 1 | ** | 5,52 | 1 | * | 0,51 | 1 | NS | 4,15 | 1 | * | 0,29 | 1 | NS |
| Temperature : Continent | 0,18 | 2 | NS | 0,07 | 2 | NS | 7,01 | 2 | * | 8,77 | 2 | * | 2,29 | 2 | NS |
| Karyotype : Sexe | | | | | | | | | | | | | 0 | 1 | NS |
| Temperature : Sexe | | | | | | | | | | | | | 4,4 | 2 | NS |
| Continent : Sexe | | | | | | | | | | | | | 2,71 | 1 | NS |
| Classic R² | 0,48 | | | 0,42 | | | 0,65 | | | 0,64 | | | 0,79 | | |
| Deviance R² | 0,45 | | | 0,42 | | | 0,6 | | | 0,61 | | | 0,87 | | |

#### Supplementary figures

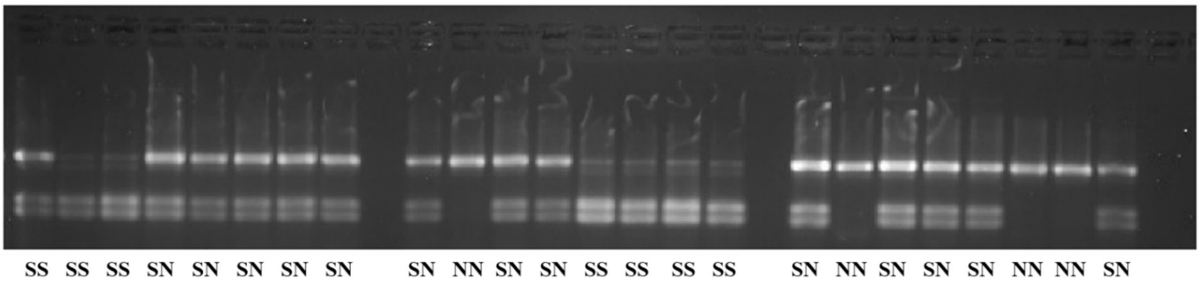

Supplementary figure 1. Electrophoresis gel performed after PCR and digestion by EcoRI to genotype inversion Cf-Inv(4.1). EcoRI cut arrangement S and not arrangement N.

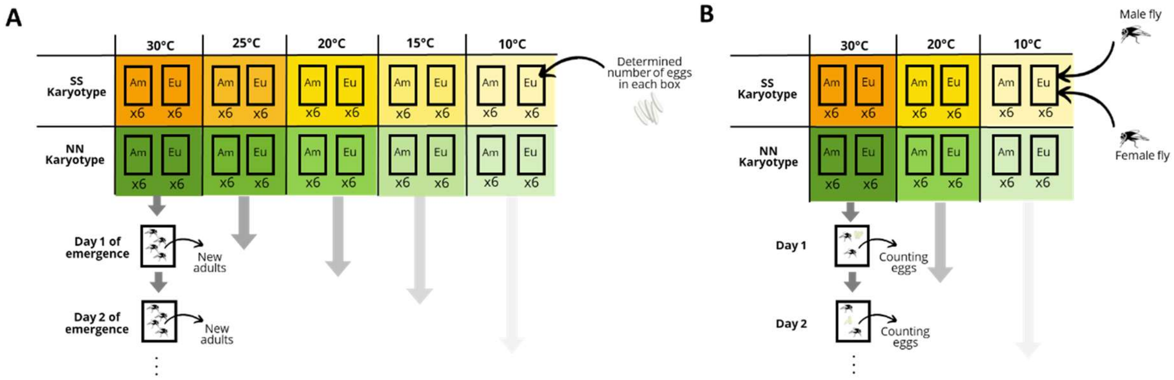

Supplementary figure 2. Experimental design for fitness measurements during development (A) and at adult stage (B).

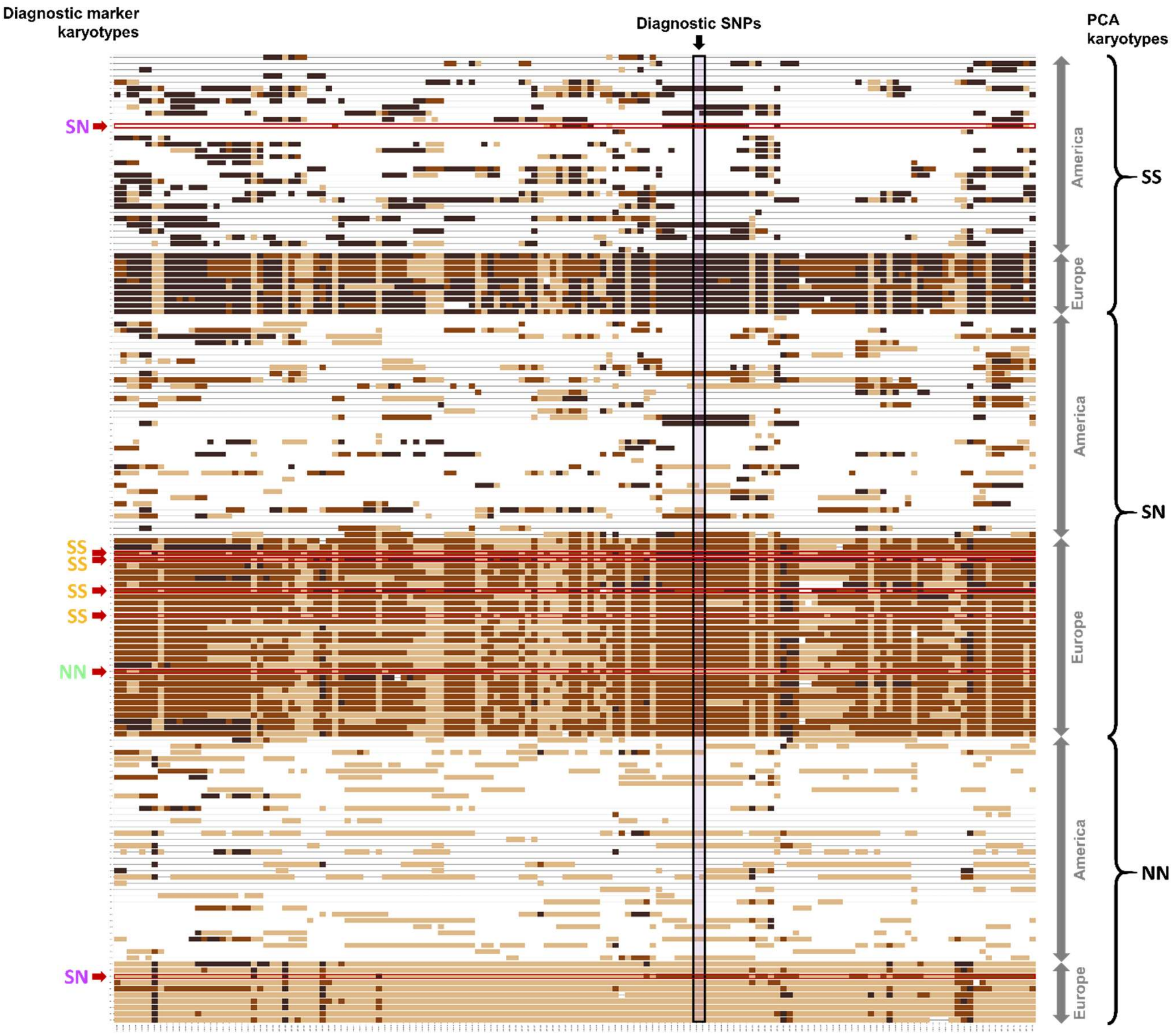

Supplementary figure 3. Genotypes at all biallelic SNPs within inversion *Cf-Inv(4.1)* between positions 1,634,841 (3 kb before diagnostic SNPs) and 1,640,843 (3 kb after diagnostic SNPs). Each line corresponds to one individual and each column to one SNP. SNPs homozygous for the reference allele are represented in beige. SNPs homozygous for the alternative allele are represented in dark brawn. Heterozygous SNPs are represented in medium brawn. The two SNPs targeted by the restriction enzyme during diagnostic assays are highlighted in purple. Individuals are grouped by inversion karyotypes as determined by the local PCA and by continents. Individuals for which the diagnostic assay gave a different karyotype than the local PCA are highlighted in red. On the left side of the graph are the karyotypes as determined by the diagnostic assay for those individuals.

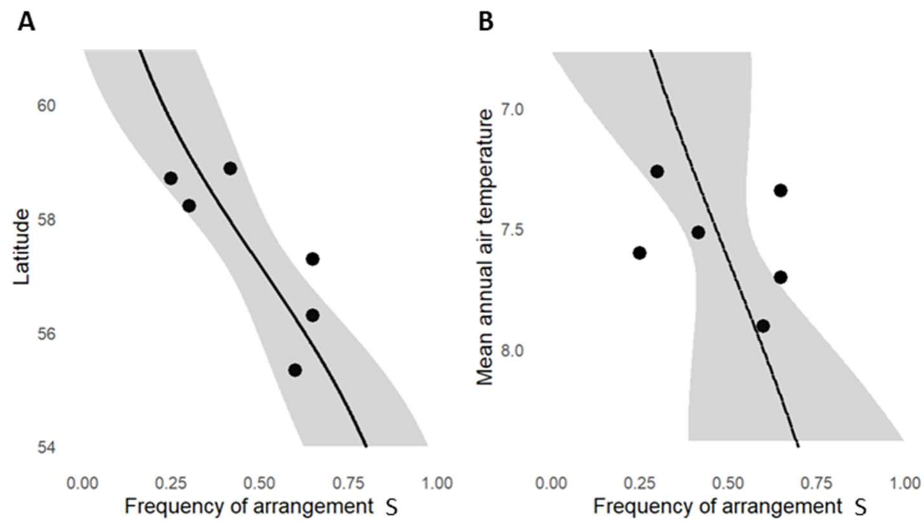

Supplementary figure 4. Clines of inversion frequency with whole-genome sequencing data. (A) Frequency of *Cf-Inv(4.1)* arrangement S as a function of latitude. (B) Frequency of *Cf-Inv(4.1)* arrangement S as a function of mean annual air temperature. A PCA was performed in the region of Cf-Inv(4.1) on whole-genome sequences of 56 individuals coming from 6 Scandinavian locations. Karyotypes were determined from the PCA. Dots represent the observed inversion frequencies in the 6 locations. Black lines are the binomial regressions adjusted on empirical data. Grey ribbons are the confident intervals 95% of the average predictions of the binomial model, which takes into account sample sizes.

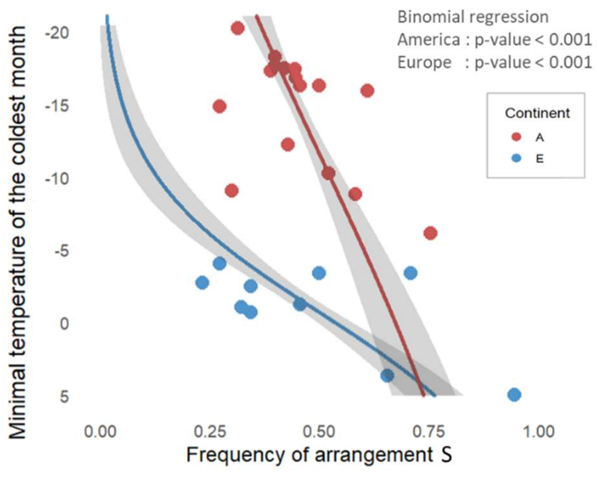

Supplementary figure 5. Inversion frequency cline along the gradient of minimal winter temperature. Frequency of *Cf-Inv(4.1)* arrangement S as a function of minimal temperature of the coldest month. Red dots represent the North American data provided by Mérot *et al.* (2021), from 16 locations sampled in September/October 2016. Blue dots represent the European data, from 10 locations sampled in March/April 2023. Red line is the binomial regression adjusted on North American data. Blue line is the binomial regression adjusted on European data. Grey ribbon is the confident interval 95% of the average predictions of the binomial model, which takes into account sample sizes.

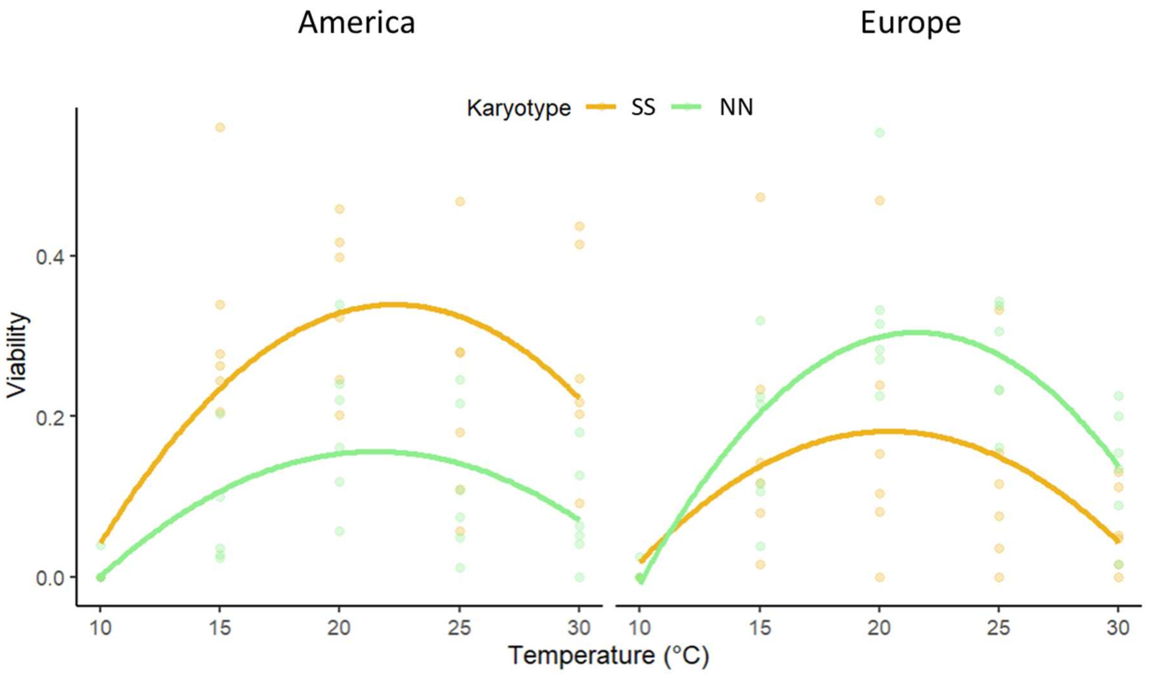

Supplementary figure 6. Thermal performance curves for viability, for each combination of karyotype and continent.

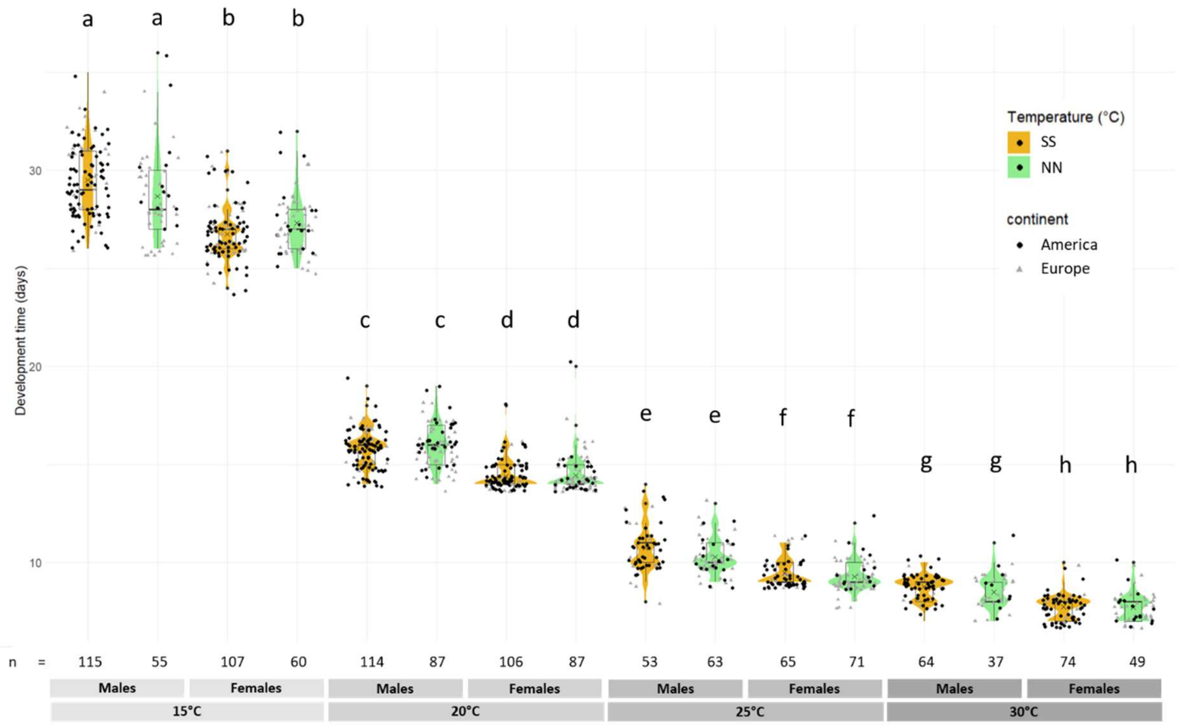

Supplementary figure 7. Development time measured as the number of days from the egg laying to the emergence of adults for each combination of temperature, karyotype, continent and sex. Data at 10°C are excluded from the analyses because the very low viability resulted in a very low sample size for development time (n = 4).

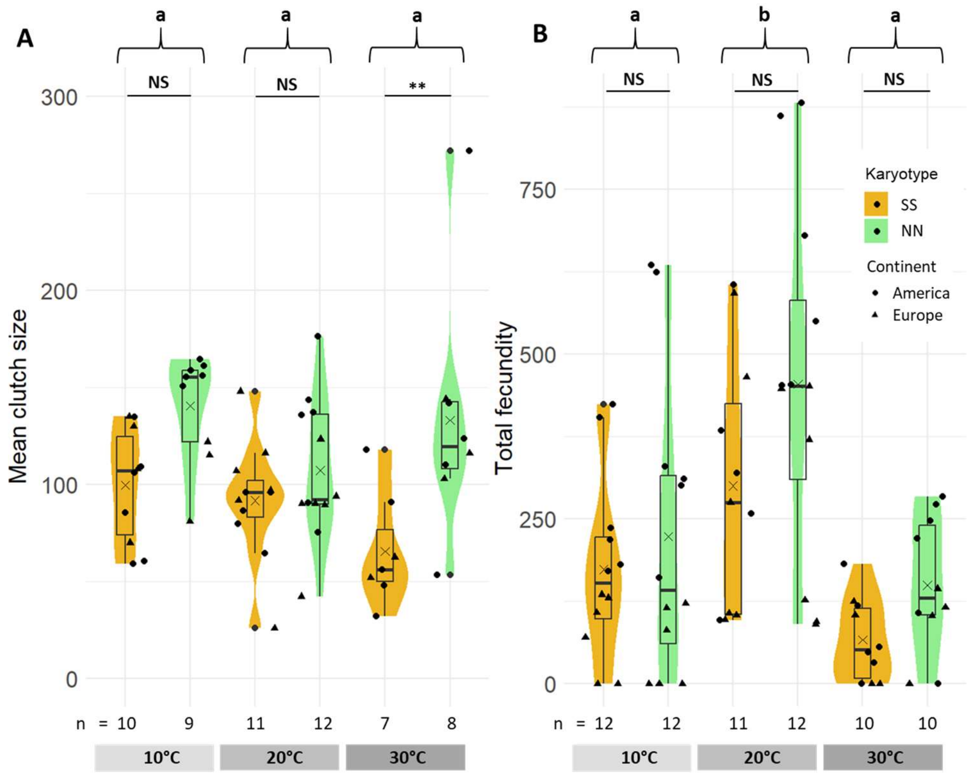

Supplementary figure 8. Mean clutch size and total fecundity. (A) Females mean clutch size for each combination of continent and karyotype. (B) Total female fecundity for each combination of temperature and karyotype.

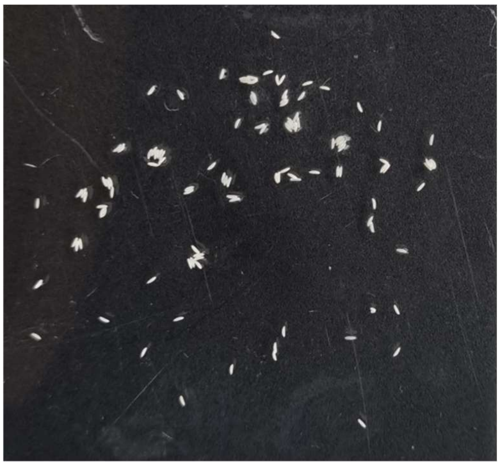

Supplementary figure 9. Picture of eggs
